# Genetic factor for twisting legume pods identified by fine-mapping of shattering-related traits in azuki bean and yard-long bean

**DOI:** 10.1101/774844

**Authors:** Yu Takahashi, Alisa Kongjaimun, Chiaki Muto, Yuki Kobayashi, Masahiko Kumagai, Hiroaki Sakai, Kazuhito Satou, Kuniko Teruya, Akino Shiroma, Makiko Shimoji, Takashi Hirano, Takehisa Isemura, Hiroki Saito, Akiko Baba-Kasai, Akito Kaga, Prakit Somta, Norihiko Tomooka, Ken Naito

## Abstract

Legumes have evolved a unique manner of seed dispersal in that the seed pods explosively split open with helical tension generated by sclerenchyma on the endocarp. During domestication, azuki bean (*Vigna angularis*) and yard-long bean (*Vigna unguiculata* cv-gr. Sesquipedalis) have reduced or lost the sclerenchyma and lost the shattering behavior of seed pods. Here we performed fine-mapping with back-crossed populations and narrowed the candidate genomic region down to 4 kbp in azuki bean and 13 kbp in yard-long bean. Among genes located in these regions, we found MYB26 genes encoded truncated proteins in both the domesticated species. We also found in azuki bean and other legumes that MYB26 is duplicated and only the duplicated copy is expressed in seed pods. Interestingly, in Arabidopsis MYB26 is single copy and is specifically expressed in anther to initiate secondary wall thickening that is required for anther dehiscence. These facts indicated that, in legumes, MYB26 has been duplicated and acquired a new role in development of pod sclerenchyma. However, pod shattering is unfavorable phenotype for harvesting and thus has been selected against by human.

## INTRODUCTION

Angiosperms have evolved various manners of seed dispersal, which is often called “shattering”. Although it is vital in nature, it is critical in agriculture. It sometimes forces farmers to lose more than 20% of their annual harvest (Philbrook and Oplinger, 1989). Thus, elucidating genetic mechanisms of shattering is important not only to understand plant evolution but also to reduce harvest loss.

Among the angiosperms, legumes have evolved a unique and sophisticated manner of shattering (Fahn and Zohary, 1955). Legume seeds are enclosed in seed pods, which explosively split open with its two valves twisting helically away from each other upon maturity (Armon et al., 2011). This mechanism is much more complicated compared to cereals, where matured seeds freely fall by abscission layer developed in pedicels (stalk of individual flower) (reviewed by Dong and Wang, 2015). *Brassica* and *Arabidopsis* are more similar to legumes in that the seeds are embedded in the siliques, that also spring open upon maturity (Spence et al., 1996). However, their siliques do not exhibit any helical shape change as legume pods do. As such, knowledge obtained from rice and *Arabidopsis* cannot be simply applied to understand the shattering of legumes.

The key of helical shape change of legume seed pods is the development of thick sclerenchyma with a bilayer structure on the endocarp (inner surface of the pod). In this bilayer structure cellulose microfibrils of the outer and inner layers run at ±45° from the longitudinal axis of seed pods (Erb et al., 2013). As the matured seed pods dry, the microfibrils shrink in perpendicular directions and generate the helical force to blow the seeds off (Erb et al., 2013). In contrast, the endocarp of *Arabidopsis*, which develops cell layers with thickened secondary walls, is inflexible (Spence et al., 1996). As the matured siliques dry, the pericarp tissues shrink but the endocarp cell layers do not, generating tension to spring open the siliques from the dehiscence zone at the valve margins (Spence et al., 1996).

However, no specific genes have been identified for the sclerenchyma formation, despite several QTL analyses and genome-wide association studies have located several loci involved in legumes’ pod shattering (Dong et al., 2014; Suanum et al., 2016; Murgia et al., 2017; Lo et al., 2018; Rau et al., 2019). In soybean, the domestication-type allele of *SHAT1-5* disturbs only dehiscence zone formation and does not affect helical shape change (Dong et al., 2014). Fiber content in seed pods, which is related to sclerenchyma formation, is reduced in common bean (Murgia et al., 2017; Rau et al., 2019; Parker et al., 2019), but the responsible gene is not cloned yet. An exception is *Pdh1* gene in soybean, which reduces helical force of the sclerenchyma when disrupted (Funatsuki et al., 2014). The plants with nonfunctional PDH1 protein still initiate sclerenchyma formation, so PDH1 seems involved in later steps in sclerenchyma development.

Contrary to soybean, azuki bean (*Vigna angularis* (Willd.) Ohwi et Ohashi) and yard-long bean (*Vigna unguiculata* (L.) Walp. cv-gr. Sesquipedalis E. Westphal) seem disrupted in initiating sclerenchyma formation. In azuki bean, an important legume crop in East Asia, the seed pods of domesticated accessions normally form dehiscence zones but the pericarps are thinner and tenderer and show little helical shape change (Isemura et al., 2007; Kaga et al., 2008). In yard-long bean, which is often cultivated in Southeast Asia, pods are very long (60-100 cm) and do not show any helical shape change at all (Kongjaimun et al., 2012). In addition, the pods are extremely tender and thus young pods are favored as vegetables (Kongjaimun et al., 2013; Suanum et al., 2016). Therefore, we consider azuki bean and yard-long bean are the best material to isolate the gene for initiating sclerenchyma formation.

Thus, in this study, we performed fine mapping to identify the responsible locus for pod shattering and pod tenderness in azuki bean and yard-long bean, respectively. Interestingly, we have previously revealed that the QTLs controlling pod shattering in azuki bean and pod tenderness in yard-long bean are co-localized around 8-10 cM on the linkage group 7 (LG7) (Kongjaimun et al., 2013), which corresponds to chr7 in azuki bean (Sakai et al., 2015) and chr5 in cowpea (Lonardi et al., 2019). To narrow down the candidate region, we developed backcrossed populations and DNA markers based on the genome sequence of azuki bean (Sakai et al., 2015), cowpea (Lonardi et al., 2019) and the wild cowpea (sequenced in this study). Our efforts identified MYB26 transcription factor as the only candidate, which indicated the evolutionary origin of the unique manner of pod shattering in legumes.

## RESULTS

### Anatomical analysis of pod sclerenchyma

To confirm that azuki bean and yard-long bean have lost or reduced sclerenchyma in seed pods, we observed cross-sections of seed pods stained with phloroglucinol-HCl and found a clear-cut difference between the wild and domesticated accessions. In the wild azuki bean and cowpea, which have shattering phenotype, thick layers of sclerenchyma (∼0.3 mm) were formed on the on the endocarps of seed pods (**Fig. 1**). However, in the domesticated azuki bean and yard-long bean, which are both non-shattering, the sclerenchyma layer was slightly formed (<0.1 mm) or not formed at all, respectively (**Fig. 1**).

**Figure 1.**
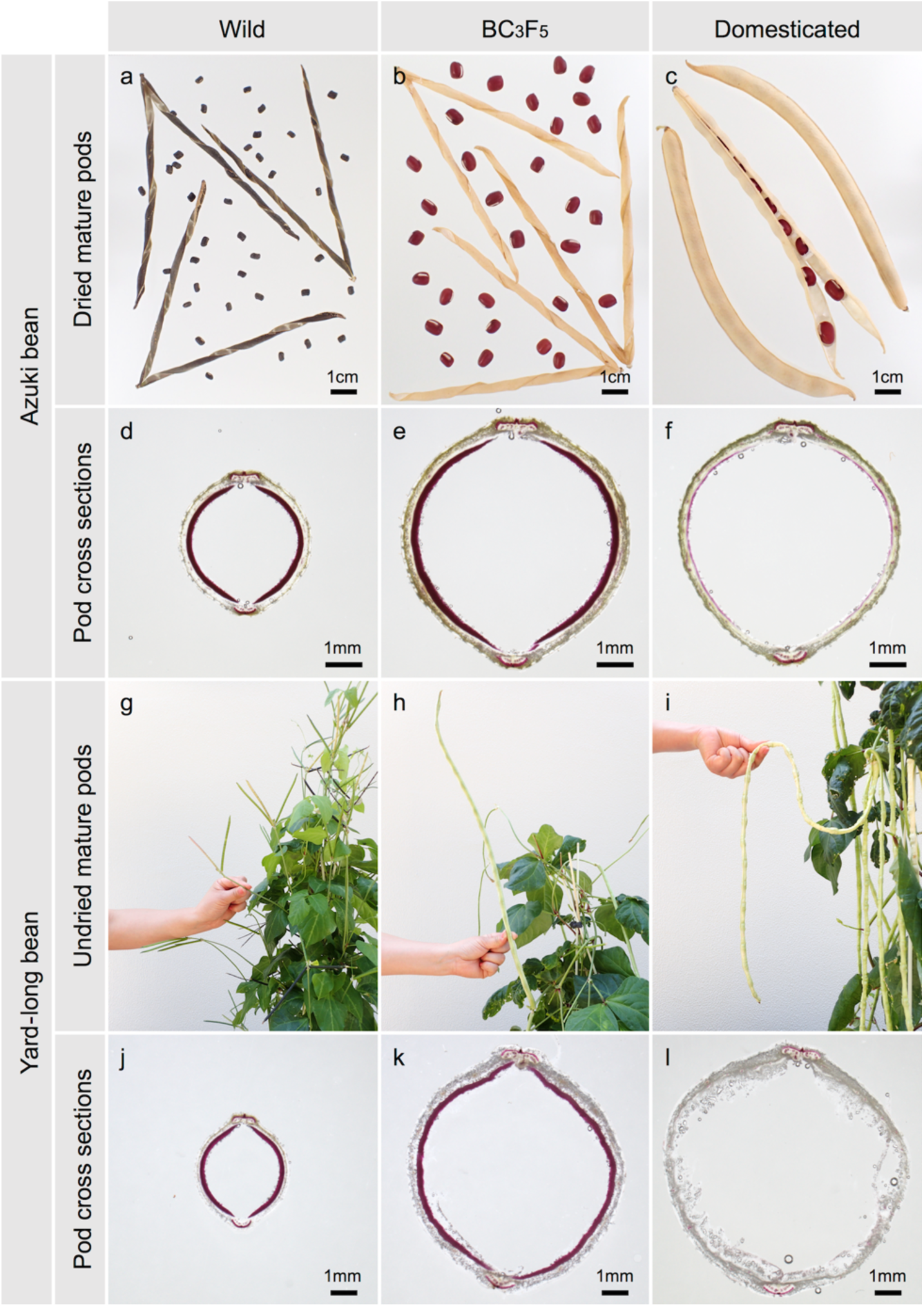
Pod phenotypes of parental accessions and BC_3_F_5_ plants with wild-type phenotypes in genetic background of domesticated genotypes. (a,d) Wild azuki bean (JP107881). (b,e) BC_3_F_5_ plant with shattering phenotype,.(c,f) Domsticated azuki bean (JP81481). (g,j) Wild cowpea (JP81610). (h,k) BC_3_F_5_ plants with hard pod phenotype. (i,l) Yard-long bean (JP89093). (a-c) Dried mature pods. (d-f, j-l) Cross sections of pods stained with phloroglucinol. (g-i) Young pods.

We also evaluated correlation between the thickness of pod sclerenchyma and helical shape change in seed pods (number of twist/cm) in the BC_3_F_5_ plants of azuki bean and yard-long bean. As a result, all the shattering plants in azuki bean population formed thicker sclerenchyma (0.15-0.20 mm) and showed stronger helical shape change (0.30-0.43) than the non-shattering plants did (0.00-0.08 mm and 0.00-0.05, respectively) (**Fig. 1, Supplementary Fig. 1**). The correlation coefficient between the sclerenchyma thickness and the helical shape change was 0.92. In the yard-long bean population, all the plants with hard pod phenotype formed sclerenchyma (0.16-0.22 mm) and showed slight helical shape change (0.09-0.12), whereas those with soft pod phenotype did not form sclerenchyma or show helical shape change at all (**Fig. 1, Supplementary Fig. 1**). The correlation coefficient between the sclerenchyma thickness and the helical shape change was 0.99.

In addition, we observed cross-sections of seed pods in a wild soybean and three domesticated soybean cultivars, and found thick sclerenchyma layers in all the accessions (**Supplementary Fig. 2**).

### Fine mapping pod shattering in azuki bean

We previously mapped the QTL for pod shattering in between CEDG182 and CEDG174 on LG7, which is around 1.4 Mbp-4.0 Mbp in Chr7 (**Fig. 2**). To more finely locate the pod shattering genetic factor, we genotyped 1,049 BC_3_F_2_ plants and selected 238 for phenotyping. The obtained phenotype and genotype data revealed the pod shattering factor was completely linked with CEDG064 and located in between SPD03 and SPD04 (**Fig. 2, Supplementary Table 2**). We selected 46 plants out of the 238 (**Supplementary Table 2**), selfed them and obtained 4,222 BC_3_F_3_ seeds.

**Figure 2.**
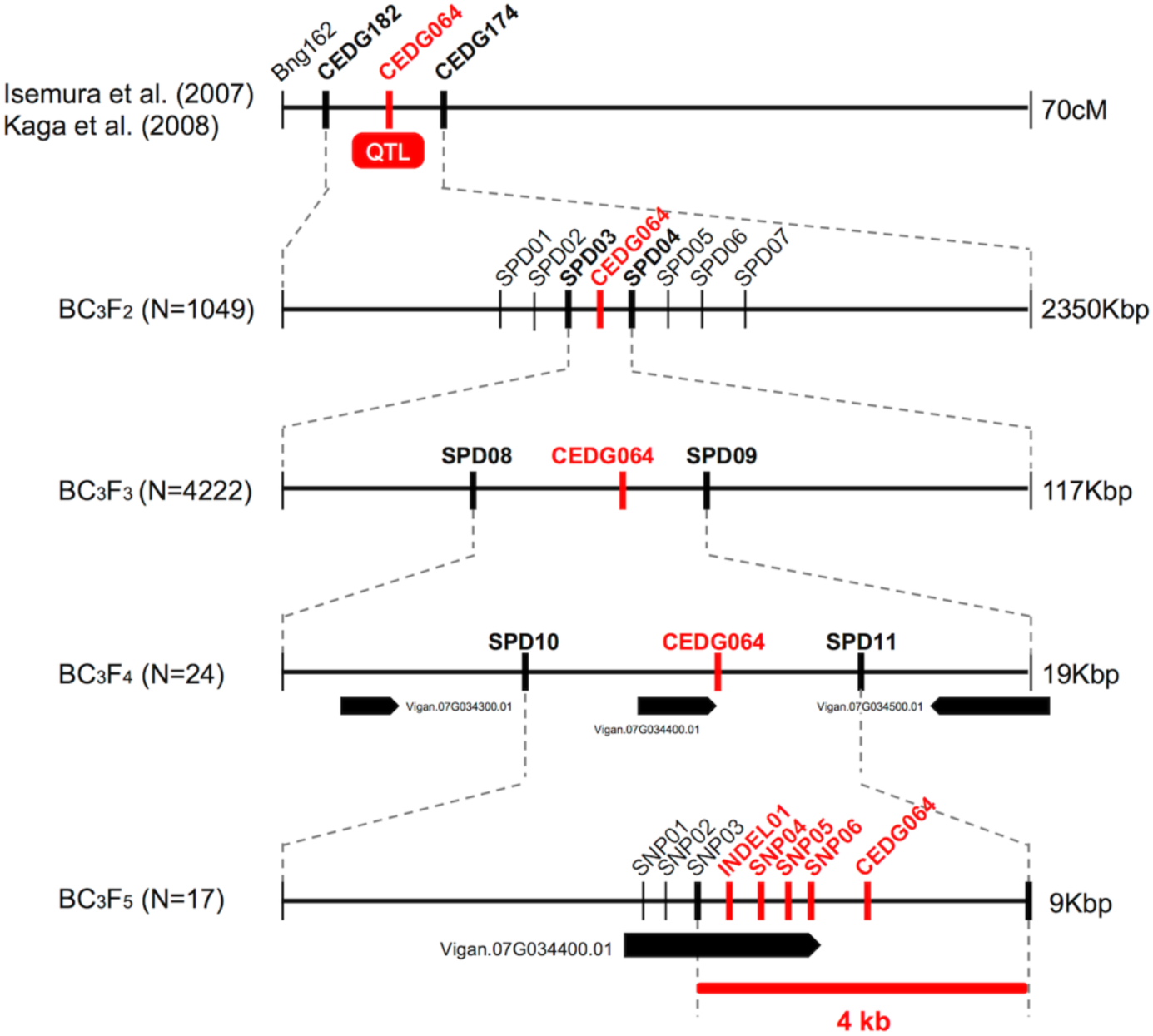
Fine mapping of pod shattering factor in azuki bean. Vertical lines indicate DNA marker sites. Red markers indicate those with complete linkage with the phenotype in the mapping population. Bold black markers indicate those neighboring the candidate region.

Of the BC_3_F_3_ seeds, we selected 53 that had recombination between SPD03 and SPD04. Phenotyping and genotyping on these 53 revealed that pod shattering factor was still completely linked with CEDG064 and was in between SPD08 and SPD09 (**Fig. 2, Supplementary Table 3**), which is about 19 kb long and contained three genes (**Fig. 2**).

Since this region seemed to have relatively higher recombination rate, we expected it would be possible to further locate the pod shattering factor. Thus, of the 53 BC_3_F_3_ plants, we selected 24 plants (see **Supplementary Table 3**) and selfed them to obtain BC_3_F_4_ seeds. We then cultivated 1-7 plants/line (81 in total, see **Supplementary Table 4**) for further genotyping and phenotyping (**Fig. 2**).

As a result, we again found CEDG064 was completely linked with the phenotype, but we also obtained 29 recombinants between SPD10 and CED064 and five between CEDG064 and SPD11 (**Supplementary Table 4**). Thus, the pod shattering factor was located in a 9 kb region between SPD10 and SPD11, where there is only one gene Vigan.07g034400 (**Fig. 2**), which encoded VaMYB26, an ortholog of AtMYB26.

To see if there were any loss-of-function mutations in VaMYB26 in azuki bean, we searched the polymorphism data between wild and domesticated azuki bean provided by *Vig*GS. The database showed that there were six SNPs (SNP01-SNP06) and one INDEL (INDEL01) within its ORF. Of these, all the SNPs were in the introns or in the 3’-UTR, but the INDEL was within the CDS (a T insertion 4 bp before the stop codon in the domesticated azuki bean). Since the wild azuki bean did not have this T insertion, the CDS could be 405 bp (125 aa) longer (**Supplementary Fig. 3**). Interestingly, a BLAST search of MYB26 gene revealed that the longer version is widely conserved across plant taxa. Thus, the VaMYB26 in the domesticated azuki bean seemed to have an immature stop codon.

To confirm the polymorphisms identified in the database, we determined the genomic and the transcribed sequences of VaMYB26 locus of both parents and found all the polymorphisms in the ORF were true (**Supplementary Fig. 3**). We also sequenced the 17 lines of BC_3_F_5_, which we obtained from the selected BC_3_F_4_ plants (those with fixed genotypes between SPD03-SPD09, see **Supplementary Table 5**), to further confirm relationship between the genotypes and the phenotype. Of the 17, one had recombination between SNP03 and INDEL01, but none had recombination between INDEL01 and CEDG064 (**Supplementary Table 5)**. Thus, the genotypes at INDEL01, SNP04-SNP06 and CEDG064 were completely linked with the pod shattering phenotype, locating the pod shattering factor within the 4 kb region between SNP03 and SPD11 (**Fig. 2, Supplementary Table 5**).

### Assembly and annotation of wild cowpea genome

We sequenced and assembled the genome of *V. unguiculata* subsp. *dekindtiana* (JP81610), which is a wild cowpea accession used to develop the mapping population of yard-long bean. We used PacBio RSII and obtained 4,413,480 raw reads with average read length of 6.0 kbp and N50 length of 10.6 kbp, covering 44.9X of the estimated size of the cowpea genome (585.8 Mbp) (Lonaridi et al., 2019) (**Supplementary Fig. 4, Supplementary Table 6**). The assembly produced 4,285 contigs covering 488.5 Mbp (83.4%) of the cowpea genome, with the contig NG50 of 438.6 kbp and the maximum contig length of 2.6 Mbp, respectively (**Supplementary Table 7**). Of the assembled sequences, 45.7% composed of transposable elements, of which LTR retrotransposons share the largest content (20.5% of the assembly) (**Supplementary Table 8**). Our *Ab initio* gene prediction detected 36,061 protein-coding genes and 27,345 of those were non-repeat related (**Supplementary Table 9**). The annotated genes of the wild cowpea contained 93.2% (85.6% complete and 7.6% partial) of the 1,440 plant genes in BUSCO v3 (Waterhouse et al., 2017).

We aligned the genome sequences of the wild cowpea to those of the reference cowpea (Lonaridi et al., 2019) and identified 5,661,319 SNPs and 1,626,169 INDELs. As expected, we detected fewer SNPS and INDELs around centromeric and pericentromeric regions but many in chromosome arms, which were presumably gene-rich regions (**Supplementary Fig. 5**).

### Fine mapping pod tenderness factor in yard-long bean

The QTL for pod tenderness was previously located between cp06388 and VR294 on LG7 (Kongjaimun et al., 2013) (**Fig. 3**), which corresponds to about 47.5-45.5 Mbp region in Chr5 of the cowpea genome (*Vigna unguiculata* v1.0, NSF, UCR, USAID, DOE-JGI, http://phytozome.jgi.doe.gov/). To more finely locate the pod tenderness factor, we designed more markers based on the SNPs and INDELs we identified above, genotyped 2,304 BC_3_F_4_ seeds and selected 195 for phenotyping (**Supplementary Table 10**).

**Figure 3.**
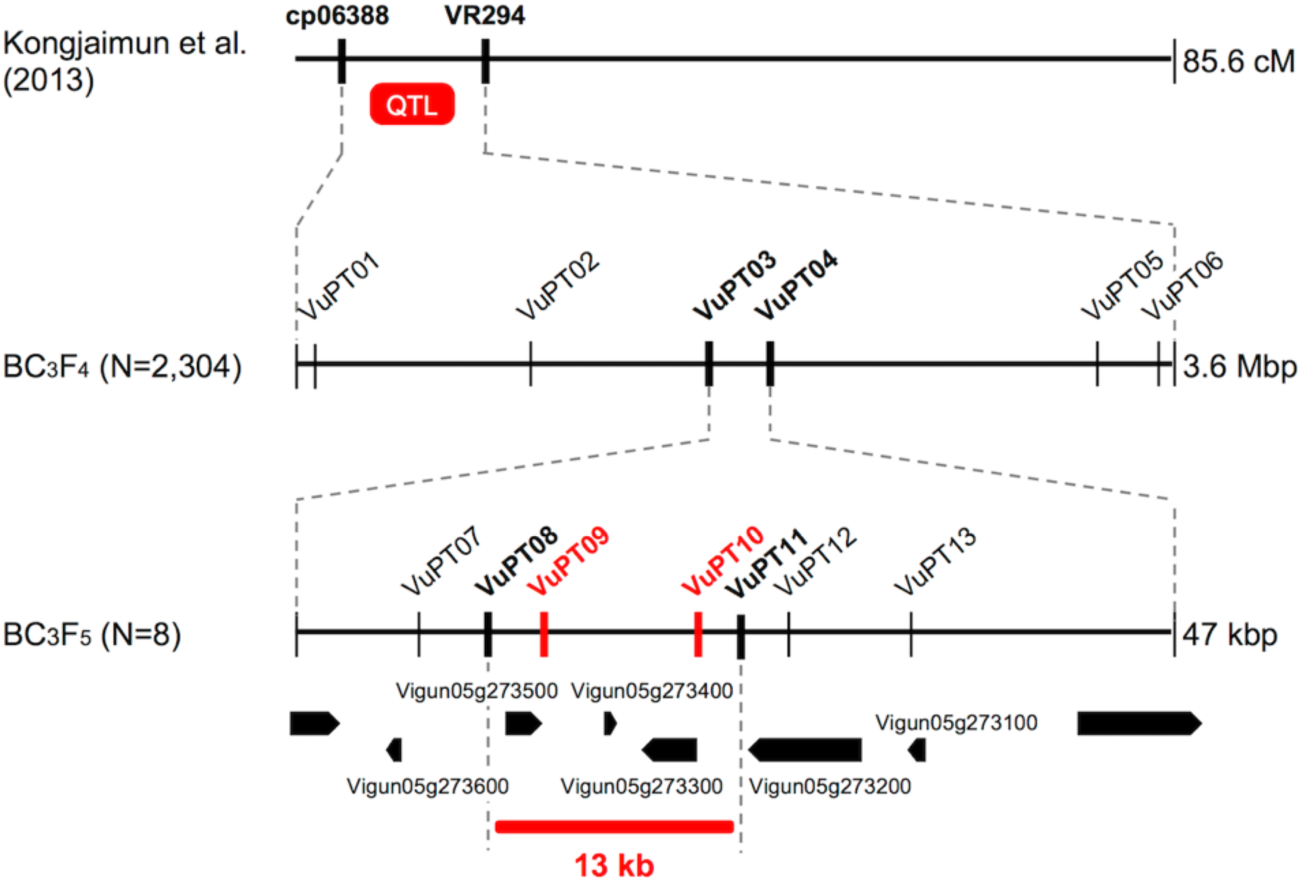
Fine mapping of pod tenderness factor in yard-long bean. Vertical lines indicate DNA marker sites. Red markers indicate those with complete linkage with the phenotype in the mapping population. Bold black markers indicate those neighboring the candidate region.

The obtained genotype and phenotype data located the pod tenderness factor between VuPT03 and VuPT04, which was about 47 kbp (**Fig. 3, Supplementary Table 10**). Although no marker was completely linked with the phenotype, 8 plants had recombination between this region (**Fig. 3, Supplementary Table 10**).

Of the 195 BC_3_F_4_ plants we tested, we selfed the 8 plants and obtained BC_3_F_5_ for further mapping. As a result, we located the candidate regions in between VuPT08 and VuPT11, which was 13 kbp long and contained 3 genes, Vigun05g27300, Vigun05g273400, and Vigun05g273500. Of note, Vigun05g273500 encoded VuMYB26 (**Fig. 3, Supplementary Table 11**).

According to the SNPs and INDELs data, Vigun05g273500 had an A to G substitution which might disrupt the junction site of the 1^st^ intron and the 2^nd^ exon (**Supplementary Fig. 6**). On the other hand, Vigun05g27300 seemed to have only synonymous SNPs, and Vigun05g273400 was a non-coding gene. All the SNPs were confirmed by directly sequencing the ORFs of these genes. Interestingly, the ORF sequences of the three genes were exactly the same between the reference cowpea and yard-long bean (**Supplementary Fig. 6**).

To test whether the A to G substitution disrupted splicing, we sequenced cDNAs of this locus prepared from the both parents. As a result, we found, in the domestication-type allele, the junction site of the 1^st^ intron and the 2^nd^ exon had shifted downstream by 8 bp, which could result in a frame-shift and an immature stop codon in the middle of the 2^nd^ exon (**Supplementary Fig. 6**). This product would encode a protein of 60 aa, which would be 305 aa shorter than that of the wild-type allele.

### Seed size increase by loss of sclerenchymal tissue

Because Murgia et al. (2017) suggested the pod shattering phenotypes load an energy cost on plants and limit seed size, we measured 100 seed weight of the mapping population (BC_3_F_2_ of azuki bean and BC_3_F_4_ of yard-long bean).

As expected, those with domestication-type trait produced larger seeds compared to those with the wild-type trait (**Fig. 4**). In the azuki bean population, the seeds of non-shattering plants were 15.4±2.4 g whereas the seeds of shattering plants were 13.3±1.9 g. In the yard-long bean population, the seeds of tender-pod plants were 13.6±1.9 g whereas the seeds of soft-pod plants were 12.9±1.7 g. The following t-test revealed that in both cases the seeds of the plants with the domestication-type phenotypes produced significantly larger seeds than the others (p<0.05).

**Figure 4.**
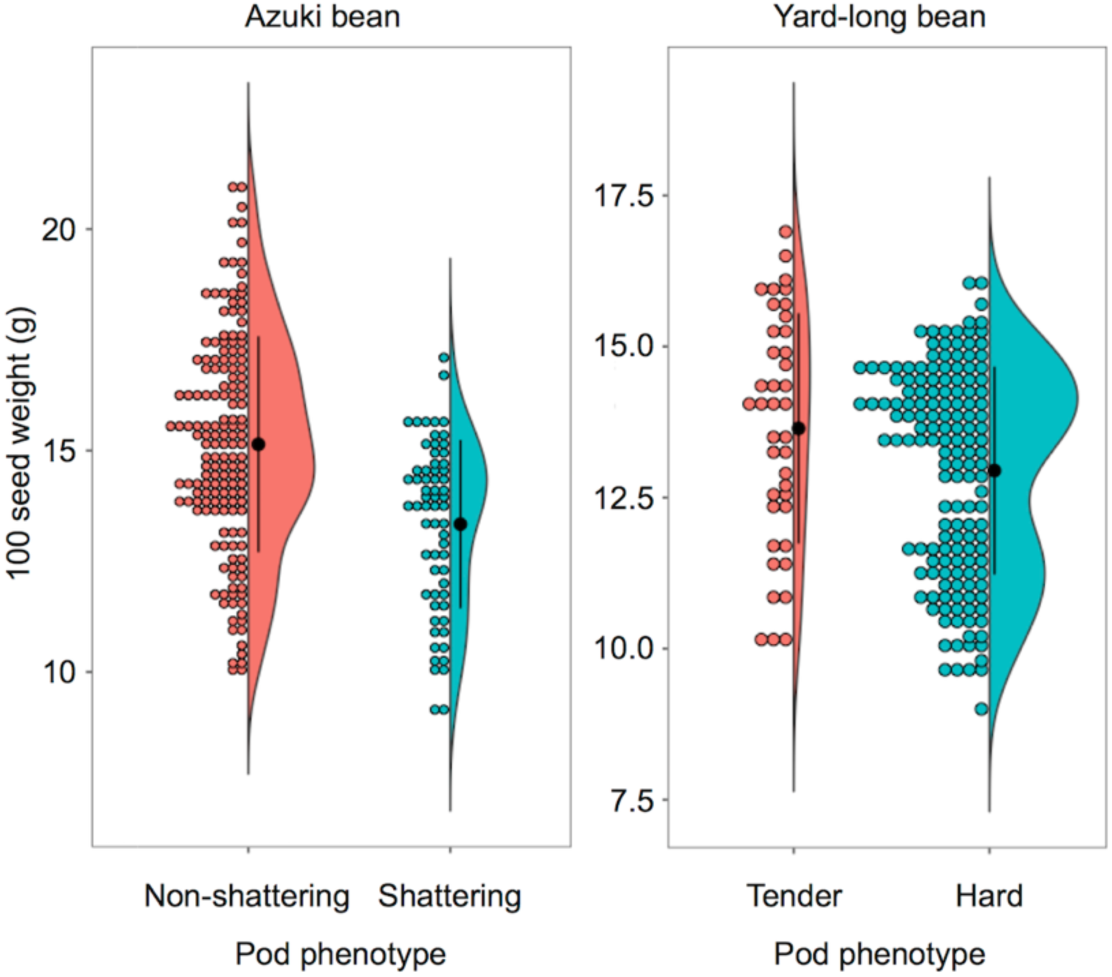
Dot-violin plot of 100 seed weight in azuki bean and yard-long bean mapping population. The bin of dot plots is 0.2 g. The black dots and vertical lines in the violin plots indicate the averages and standard deviations, respectively.

We also measured 100 seed weight of BC_3_F_5_ plants and observed the same trend (**Supplementary Fig. 7)**. In azuki bean, the seeds of non-shattering plants were 9.7±1.4 g whereas the seeds of shattering plants were 8.7±1.2 g (p<0.05). In yard-long bean, although not significant, the seeds of tender-pod plants were 14.2±1.5 g whereas the seeds of hard-pod plants were 12.2±1.2 g.

### Phylogenetic analyses of MYB26

To elucidate evolutionary origin of VaMYB26 and VuMYB26, we reconstructed a phylogenetic tree of MYB26 genes of azuki bean, cowpea, common bean, soybean and *Arabidopsis* using AtMYB55 as outgroup (**Fig. 5**). Before reconstructing phylogenetic tree, we BLASTed the VaMYB26 sequence and found one or more paralogous copies in all the legume genomes investigated. We designated them as MYB26a and MYB26b (see materials and methods). The reconstructed phylogenetic tree formed a single clade of MYB26, with subclades of MYB26a and MYB26b (**Fig. 5**).

**Figure 5.**
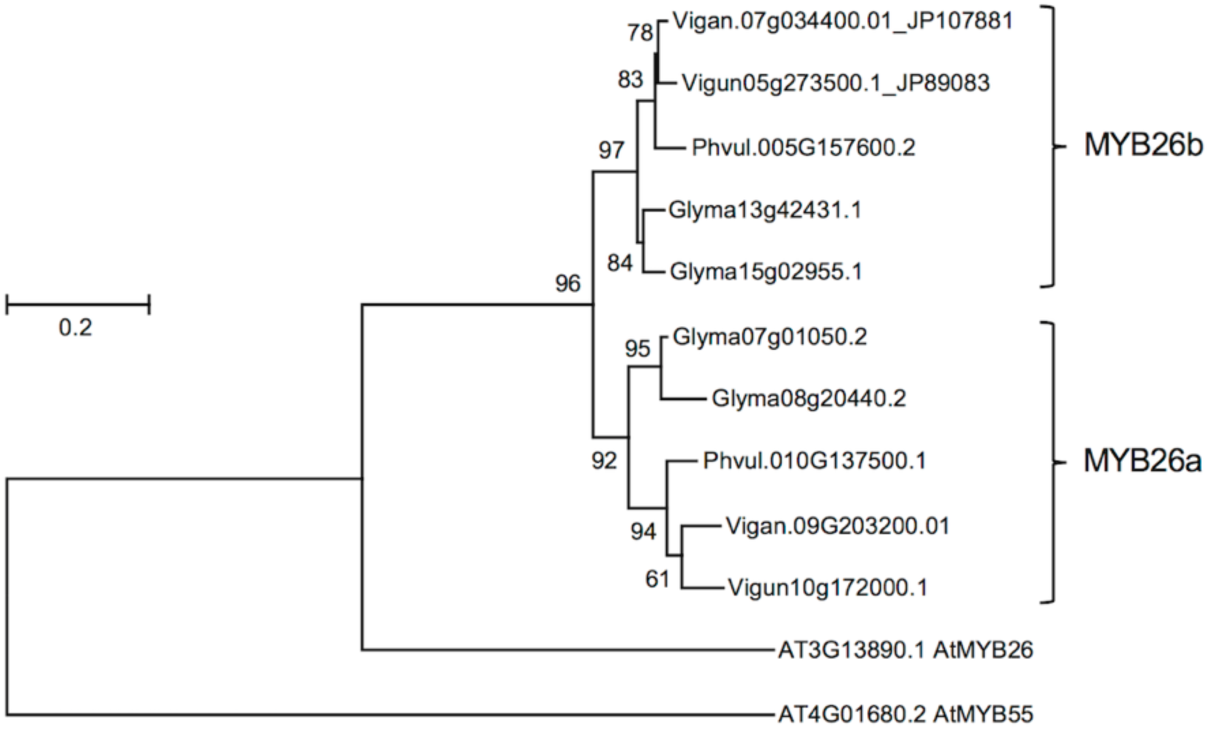
Phylogenetic tree of MYB26 by Maximum Likelihood method based on the JTT matrix-based model. The tree is reconstructed to scale, with branch lengths measured in the number of substitutions per site. The analysis involved 12 amino acid sequences. All positions containing gaps and missing data were eliminated.

### Expression profiles of MYB26

Because AtMYB26 is hardly transcribed in the siliques (https://www.arabidopsis.org/servlets/TairObject?type=locus&name=At3G13890), we examined whether the MYB26 genes were transcribed in legume pods by extracting their expression profiles from the databases of azuki bean, common bean and soybean. As a result, we found MYB26b is transcribed in the pods of all the legume species but MYB26a is not (**Supplementary Fig. 7**). In azuki bean, VaMYB26a is transcribed in leaf, stem and flowers whereas VaMYB26b is transcribed in stem, root, flower and pod. In common bean, PvMYB26a is transcribed in flower, and PvMYB26b is transcribed in stem and pod. In soybean, GmMYB26a1 is transcribed in root and pod, and other copies are transcribed only in pod. However, GmMYB26b1 is by far the most transcribed copy in pod.

## DISCUSSION

In this study, we identified VaMYB26b (**Fig. 2**) and VuMYB26b (**Fig. 3**) are the key factors for sclerenchyma formation on the endocarp of seed pods in azuki bean and yard-long bean (**Fig. 1**). Both the loci encode truncated proteins due to immature stop codons (**Supplementary Figs. 3, 6**), leading to reduction or loss of sclerenchyma, which is required to generate the helical tension for splitting open legume pods (**Fig. 1, Supplementary Fig. 1**). Thus, the seed pods of azuki bean have lost shattering ability (Isemura et al., 2007) and those of yard-long bean have acquired tenderness and become edible (Kongjaimun et al. 2013, Suanum et al. 2016). Though recent genetic and genomic studies on common bean have identified several candidate loci for pod shattering, PvMYB26b is not attributed to the phenotype so far (Rau et al., 2019; Parker et al., 2019). Of the candidate loci, qPD5.1-*Pv* is the locus harboring PvMYB26b, but Rau et al. (2018) found the SNPs in this gene was not best associated with the trait. In contrast, we narrowed the candidate region down to 4 kbp in azuki bean, where only VaMYB26b is present, and 13 kbp in yard-long bean, where three genes are present including VuMYB26b. The domestication-type alleles of VaMYB26b and VuMYB26b encode truncated proteins, which are likely non- or partially-functional.

We have to admit that availability of whole genome sequences greatly facilitated this study. Although the lower coverage and shorter read length of our assembly resulted in ∼10 times lower contiguity than the reference cowpea genome (Lonardi et al., 2019), its accuracy was good enough as all the SNPs between the two assembly, which we tried to use as markers, were confirmed to be true (**Supplementary Tables 5,11, Supplementary Figs 3,6**). We also re-emphasize the power of map-base cloning, which is a classic, time-consuming but reliable approach to isolate responsible genes.

Thanks to the whole genome sequences, our results also indicated an evolutionary pathway of the unique phenotype of pod shattering in legumes. In *Arabidopsis*, AtMYB26 is a single copy gene which is strongly expressed in the anther and promote secondary wall thickening in the endothelium (Yang et al., 2007). Interestingly, this secondary wall thickening is critical for anther dehiscence and thus for releasing the matured pollen (Yang et al., 2007). However, this gene is hardly expressed in siliques and is not involved in pod dehiscence (Yang et al., 2007). In contrast, MYB26 has undergone ancestral duplication in legumes (**Fig. 5**), and one of the paralogs (MYB26b) has become transcribed in the seed pods (**Supplementary Fig. 8**). Thus, MYB26b has acquired a new role in secondary wall thickening in seed pods, leading to development of sclerenchyma with helical force that enables the pods to explosively split open. We consider this is a good example of neofunctionalization, where a duplicated copy acquires increased expression in one or more tissues/organs (Duarte et al., 2006).

We also consider the knowledge obtained in this study is applicable to improve soybean or other legume crops. As described before, soybean has reduced shattering ability by disturbing dehiscence zone formation between the valves (Dong et al., 2014) and by reducing helical tension of the sclerenchyma (Funatsuki et al., 2014). However, it still forms thick sclerenchyma in the seed pods (**Supplementary Fig. 2**), which causes shattering problems especially under drought condition (Philbrook and Oplinger, 1989). Thus, loss-of-function mutations or genome-editing in GmMYB26s may reduce sclerenchyma and enforce resistance to shattering. In addition, our findings indicate that loss or reduction of sclerenchyma may result in a slight (∼10%) increase in seed size (**Fig. 4, Supplementary Fig. 7**). Murgia et al. (2017) also reported that common bean plants with non-shattering genotypes produce slightly larger seeds than those with shattering genotypes Thus, our results supported their idea that pod shattering loads an energy cost for plants and suppression of sclerenchyma formation could save carbon source that could be allocated for seed production (Murgia et al., 2017). As such, MYB26 may serve not only for reducing harvest loss but also for directly increasing seed size.

Last but not least, our results indicated the domestication-type allele of VuMYB26 has been selected during the primary domestication of cowpea in Africa (Lush and Evans, 1981), given the reference cowpea and yard-long bean shared exactly the same sequences in this locus (Supplementary Fig. 6). Although the domestication-type allele of VuMYB26b contributes the most part of pod tenderness in yard-long bean, it should have been originally selected for non-shattering trait of cowpea. Indeed, the QTL analysis on pod tenderness (Kongjaimun et al., 2013) and pod fiber content (Suanum et al., 2016) identified one major QTL, which overlaps the VuMYB26 locus, and other minor QTLs. Thus, the complete loss of pod sclerenchyma in yard-long bean required a few more genetic alterations that arose during the secondary domestication in Asia (Faris, 1965).

In conclusion, our map-based cloning approach with support of whole genome sequences identified MYB26b as an only candidate for development of pod sclerenchyma, which generates helical tension of legumes’ shattering pods. Our findings suggest MYB26b might be a typical case of duplication-neofunctionalization theory, and a good target to improve resistance to pod shattering in soybean and other legumes.

## EXPERIMENTAL PROCEDURES

### Plant materials and growth condition

All the accessions used in this study were provided by NARO genebank (Tsukuba, Japan) (Fig. 1, Supplementary Fig. 1).

For azuki bean, we started from the BC_1_F_2_ plants derived from a cross between domesticated azuki bean (JP81481, the recurrent parent) and a wild relative, *Vigna nepalensis* Tateishi & Maxted (JP107881) (Isemura et al., 2007). Of them, we selected one BC_1_F_2_ plant where the locus for pod shattering is fixed with the wild type allele but most of other loci are fixed with domestication-type alleles, and crossed again to the recurrent parent. We further backcrossed the obtained BC_2_F_1_ plants to obtain BC_3_F_1_ plants. We selfed them and obtained BC_3_F_2_ seeds from those with pod shattering phenotype. For BC_3_F_3_, BC_3_F_4_ and BC_3_F_5_ populations, we kept selecting and selfing those with recombination within the candidate region. We also included plants that were heterozygous throughout the candidate region for obtaining more recombination, and some that are homozygous in the same region to reconfirm the relationship of genotype and the phenotype.

For yard-long bean, we started from BC_1_F_2_ plants derived from a cross between yard-long bean (JP89083, the recurrent parent) and a wild cowpea, *Vigna unguiculata* subsp. *dekindtiana* (Harms) Verdc. (JP81610) (Kongjaimun et al., 2013). Of them, we selected one BC_1_F_2_ plant that are fixed with wild type allele at the locus for pod tenderness but are fixed with domestication-type at most other loci. The selected BC_1_F_2_ plants were backcrossed to the recurrent parent, and further backcrossed to obtain BC_3_F_1_ plants. We selfed them and obtained BC_3_F_2_ seeds from those with hard pods phenotype. We kept selecting and selfing up to BC_3_F_5_ populations as done in azuki bean.

All the plants were grown from July through November in a field located in Tsukuba city, Japan except BC_3_F_5_ plants of yard-long bean, that were grown in a greenhouse from September through January.

### Observation of sclerenchymal tissue

We observed sclerenchymal tissues in seed pods by staining lignin with the Wiesner (phloroglucinol-HCl) reaction (Pormar et al., 2002). We harvested three undried mature seed pods per plant, sliced them to 200 μm sections with a Plant Microtome MTH-1 (Nippon Medical & Chemical Instruments Co., Ltd. Osaka, Japan), incubated in phloroglucinol-HCL solution for 1 min and observed with stereoscopic microscope SZX7 (OLYMPUS, Tokyo) for BC_3_F_5_ plants of azuki bean and yard-long bean. Phloroglucinol-HCL solution was prepared by dissolving 1 g of phloroglucin in 50 ml ethanol and adding 25 ml of concentrated hydrochloric acid.

### Phenotyping

For azuki bean populations, we evaluated pod shattering by counting number of twist in pod, as described in Isemura et al. (2007). For BC_3_F_2_ and BC_3_F_3_, we harvested five pods per plant before shattering and measured the length. We then incubated the pods at 60°C to let them totally dry. Unless the pods dehisced, we manually opened the pods and counted the number of twists and calculated the number of twists/cm. For BC_3_F_4_ and BC_3_F_5_, we harvested 10 pods per plant, let them dry and evaluated the rate of shattering pods before counting the number of twists.

For yard-long bean, we manually evaluated the tenderness of five young pods per plant by binarizing hard or soft.

We also measured 100 seed weight on BC_3_F_2_ and BC_3_F_5_ plants of the azuki bean population and BC_3_F_4_ and BC_3_F_5_ plants of the yard-long bean population.

### Sequencing and assembly of the wild cowpea genome

We sequenced the whole genome of wild cowpea (JP81610) with RSII sequencer (Pacific Biosciences, Menlo Park, CA), as we have done previously for azuki bean (Sakai et al., 2015) DNA was isolated from 1 g of unexpanded leaves with CTAB method and purified with Genomic Tip 20/G (Qiagen K. K. Tokyo). The extracted DNA was sheared into 20 kb fragments using g-TUBE (Covaris, MA, USA) and converted into 20 kb SMRTbell template libraries. The library was size selected for a lower cutoff of 10 kb with BluePippin (Sage Science, MA, USA). Sequencing was performed on the PacBio RS II using P6 polymerase binding and C4 sequencing kits with 360 min acquisition. In total, 21 SMRT cells were used to obtain ∼26.4 Gb of raw reads. Raw sequence data generated in this study are available at DDBJ (https://www.ddbj.nig.ac.jp/index.html) under the BioProject ID PRJDB8129.

In total, 4 million PacBio reads were used for *de novo* assembly with Canu v1.6 under the default settings (corOvlErrorRate = 0.2400, obtOvlErrorRate = 0.0450, utgOvlErrorRate = 0.0450, corErrorRate = 0.3000, obtErrorRate = 0.0450, utgErrorRate = 0.0450, cnsErrorRate = 0.0750). About 23.5x error corrected and trimmed reads longer than 1,000 bp were assembled to contigs. Assembled contigs were polished by PacBio raw reads by using arrow in GenomicConsensus v2.3.2 (Pacific Biosciences of California, Inc.).

Repeat detection was conducted by RepeatMasker ver 4.0 (http://repeatmasker.org). A *de novo* repeats library of wild cowpea genome constructed by RepeatModeler ver 1.0.11 (http://www.repeatmasker.org) and the Fabaceae repeats library in RepBase24.02 (https://www.girinst.org) were used for the prediction.

*Ab initio* gene prediction was done with AUGUSTUS (version 3.3.2) (Stanke et al., 2008). A set of gene annotation information of recently published cowpea genome was used for training AUGUSTUS (Lonardi et al., 2019). We trained a new model twice using 1,000 high-confidential genes selected based on abundance of annotations of domains, pathway networks and Gene ontology information. BUSCO v3 (Waterhouse et al., 2017) was used to evaluate protein sequences of annotated genes.

SNPs and short indels were detected by using MUMmer v3.23 (Kurtz et al., 2004). Genome alignment of our wild cowpea assembly and the reference cowpea genome (Lonardi et al., 2019) was conducted by the nucmer command with the following options: --maxgap=500 --mincluster=100. Then one to one alignment was extracted with delta-filter command with an option: −1. Based on the alignments, SNPs and indels were reported by the show-snps command.

### Genotyping

We extracted DNA from the seeds as described by Kamiya and Kiguchi (2003), and genotyped them by fragment analysis with capillary electrophoresis for SSRs (simple sequence repeats) and INDELs (insertion/deletion) as described by Isemura et al. (2007) or by directly sequencing SNP (single nucleotide polymorphisms) sites (see “direct sequencing” below). The information of markers we used are summarized in **Supplementary Table 1**.

We designed primers by Primer3 (Untergasser et al., 2012) to amplify polymorphic sites between the domesticated azuki bean and *V. nepalensis* that are available in *Vigna* Genome Server (*Vig*GS, https://viggs.affrc.go.jp, Sakai et al., 2016), and those between the domesticated cowpea genome in Legume Information System (https://legumeinfo.org/) and the wild cowpea genome sequenced in this study. Parameters for designing primers on Primer3 (http://primer3.ut.ee) were 20-30 bp in length, 55-65°C in annealing temperature and 100-350 bp and >700 bp in expected length of amplified fragments for fragment analysis and direct sequencing, respectively.

For azuki bean, the markers we used were as follows:

> BC_3_F_2_: CEDG064 and SPD01-SPD07
>
> BC_3_F_3_: CEDG064, SPD03, SPD04, SPD08, SPD09
>
> BC_3_F_4_: CEDG064 and SPD08-SPD11
>
> BC_3_F_5_: CEDG064, SPD10, SPD11 and the sequencing primers for Vigan.07G034400

For yard-long bean, the markers we used were as follows:

> BC_3_F_4_: VuPT01-VuPT06
>
> BC_3_F_5_: VuPT03, VuPT04, VuPT07-VuPT13 and the sequencing primers for Vigun05g273300, Vigun05g273400 and Vigun05g0723500.

### Direct sequencing

To sequence the (potentially) polymorphic sites, we amplified the template DNA with AmpliTaq Gold 360 Master Mix (Thermo Fisher Scientific K. K., Tokyo), performed sequencing reaction with BigDye Terminator v3.1 (Thermo Fisher Scientific K. K., Tokyo), and sequenced with ABI Genetic Analyzer 3130xl (Thermo Fisher Scientific K. K., Tokyo), according to the provider’s protocol.

### Determining transcribed sequences of MYB26b

To sequence both the domesticated and the wild alleles of MYB26a gene, we sequenced the transcribed sequences of Vigan.07g034400.01 and Vigun05g273300.01. We extracted total RNA from 100 mg of flower buds right before flowering of all the parental accessions using RNeasy Plant Mini Kit (QIAGEN) with RNase-free DNase I (Invitrogen). Total RNA of 1 μg was converted into first-strand cDNA with Super Script II Reverse Transcriptase (Invitrogen) and Oligo(dT)12-18 Primer (Invitrogen) following the manufacturer’s instructions. The cDNA sequences of MYB26a (Vigan.07g034400.01 and Vigun05g273300.01) were then amplified and sequenced with the primers (Supplementary Table 1), and then transferred to the direct-sequencing protocol described above.

### Phylogenetic analyses

For phylogenetic analysis we used amino the acid (aa) sequence of Vigan.07g034400.01 (VaMYB26a) as a query and retrieved its orthologs and paralogs from azuki bean, cowpea, common bean, soybean, and Arabidopsis from *Vig*GS (http://viggs.dna.affrc.go.jp/) (Sakai et al., 2015b), Legume Information System (https://legumeinfo.org/) (Dash et al., 2015) and TAIR (https://www.arabidopsis.org/). The retrieved sequences were: Vigan.09g203200.01(VaMYB26b), Vigun05g273500.1 (VuMYB26a), Vigun10g172000.1 (VaMYB26b), Phvul.005G157600.2 (PvMYB26a), Phvul.010G137500.1 (PvMYB26b), Glyma13g42431.1 (GmMYB26a1), Glyma15g02955.1 (GmMYB26a2), Glyma07g01050.2 (GmMYB26b1), Glyma08g20440.2 (GmMYB26b2), and AT3G13890.1 (AtMYB26). However, we replaced the aa sequences of VaMYB26a and VuMYB26a by those isolated from the wild azuki bean and the wild cowpea, respectively, because the sequences in the database were domestication-type alleles with immature stop codons. We also included a paralog of AtMYB26, AT4G01680.2 (AtMYB55), as outgroup. We aligned the obtained sequences to each other by Clustal W, removed the gaps and reconstructed a phylogenetic tree by a maximum likelihood method based on the JTT model (Jones et al., 1992) in MEGA 7.0 (Kumar et al., 2016).

### Expression profiles

We extracted expression profiles of MYB26 genes of azuki bean, common bean and soybean from *Vig*GS, the *Phaseolus vulgaris* Gene Expression Atlas (*Pv*GEA) (http://plantgrn.noble.org/PvGEA/index.jsp) and Soybean eFP Browser (http://bar.utoronto.ca/efpsoybean/cgi-bin/efpWeb.cgi).

## ACKNOWLEDGEMENTS

This work was supported by JSPS KAKENHI Grant Number 13J09808 and 26850006. It was also partially supported by Research Supporting Program of the Advanced Analysis Center, National Agriculture and Food Research Organization (NARO) and the Genebank Project, NARO.

## AUTHOR CONTRIBUTIONS

YT, PS, AK, NY and KN, planned the study.

YT, AK, CM, YK, TI, HS, AK, PD, TN and KN performed experiments and collected data.

MK, HS, KS, KT, AS, MS and TH performed genome sequencing, *de novo* assembly and gene annotation.

YT, AK, HS, MK and KN analyzed the data. YT, AK, TN and KN wrote the paper.

## REFERENCES

Armon, S., Efrati, E., Kupferman, R. and Sharon, E. (2011) Geometry and mechanics in the opening of chiral seed pods. Science, 333, 1726–1730.

Dash, S., Campbell, JD., Cannon, E.K., Cleary, A.M., Huang, W., Kalberer, S.R., Karingula, V., Rice, A.G., Singh, J., Umale, P.E., Weeks, N.T., Wilkey, A.P., Farmer, A.D. and Cannon, S.B. (2016) Legume information system (LegumeInfo.org): a key component of a set of federated data resources for the legume family.. Nucleic Acids Research, 44, D1181–D1188.

Di Vittori, V., Gioia, T., Rodriguez, M., Bellucci, E., Bitocchi, E., Nanni, L., Attene, G., Rau, D. and Papa, R. (2019) Convergent Evolution of the Seed Shattering Trait. Genes,10, 68.

Dong, Y., Yang, X., Liu, J., Wang, B.H., Liu, B.L. and Wang, Y.Z. (2014) Pod shattering resistance associated with domestication is mediated by a NAC gene in soybean. Nature Communications, 5, 3352.

Dong, Y. and Wang, Y.Z. (2015) Seed shattering: from models to crops. Frontiers in Plant Science, 6, 476.

Duarte, J.M., Cui, L., Wall, P.K., Zhang, Q., Zhang, X., Leebens-Mack, J., Ma, H., Altman, N. and dePamphilis, C.W. (2006) Expression pattern shifts following duplication indicative of subfunctionalization and neofunctionalization in regulatory genes of Arabidopsis. Molecular Biology and Evolution, 23, 469–478.

Erb, R.M., Sander, J.S., Grisch, R. and Studart, A.R. (2013) Self-shaping composites with programmable bioinspired microstructures. Nature Communications, 4, 1712.

Fahn, A. and Zohary, M. (1955) On the pericardial structure of the legumen, its evolution and relation to dehiscence. Phytomorphology, 5, 99–111.

Faris, D.G. (1965) The origin and evolution of the cultivated forms of *Vigna sinensis*. Canadian Journal of Genetics and Cytology, 7, 433–452.

Funatsuki, H., Suzuki, M., Hirose, A., Inaba, H., Yamada, T., Hajika, M., Komatsu, K., Katayama, T., Sayama, T., Ishimoto, M. and Fujino, K. (2014) Molecular basis of a shattering resistance boosting global dissemination of soybean. Proceedings of the National Academy of Sciences, USA, 111, 17797–17802.

Isemura, T., Kaga, A., Konishi, S., Ando, T., Tomooka, N., Han, O.K. and Vaughan, D.A. (2007) Genome dissection of traits related to domestication in azuki bean (*Vigna angularis*) and comparison with other warm-season legumes. Annals of Botany, 100, 1053–1071.

Jones, D.T., Taylor, W.R. and Thornton, J.M. (1992) The rapid generation of mutation data matrices from protein sequences. Bioinformatics, 8, 275–282.

Kaga, A., Isemura, T., Tomooka, N. and Vaughan, D.A. (2008) The genetics of domestication of the azuki bean (*Vigna angularis*). Genetics, 178, 1013–1036.

Kamiya, M. and Kiguchi, T. (2003) Rapid DNA extraction method from soybean seeds. Breeding Science, 53, 277–279.

Kongjaimun, A., Kaga, A., Tomooka, N., Somta, P., Vaughan, D.A. and Srinives, P. (2012) The genetics of domestication of yardlong bean, Vigna unguiculata (L.) Walp. ssp. unguiculata cv.-gr. Sesquipedalis. Annals of Botany, 109, 1185–1200.

Kongjaimun, A., Somta, P., Tomooka, N., Kaga, A., Vaughan, D.A. and Srinives, P. (2013) QTL mapping of pod tenderness and total soluble solid in yardlong bean [*Vigna unguiculata* (L.) Walp. subsp. unguiculata cv.-gr. sesquipedalis]. Euphytica, 189, 217.

Konishi, S., Izawa, T., Lin, S.Y., Ebana, K., Fukuta, Y., Sasaki, T. and Yano, M. (2006) An SNP caused loss of seed shattering during rice domestication. Science, 312, 1392–1396.

Kumar, S., Stecher, G. and Tamura, K. (2016) MEGA7: Molecular Evolutionary Genetics Analysis Version 7.0 for Bigger Datasets. Molecular Biology and Evolution, 33, 1870–1874.

Kurtz, S., Pillippy, A., Delcher, A.L., Smoot, M., Shumway, M., Sntonescu, C. and Salzberg, S.L. (2004) Versatile and open software for comparing large genomes. Genome Biology, 5, R12.

Lo, S., Muñoz-Amatriaín, M., Boukar, O., Herniter, I., Cisse, N., Guo, Y.N., Roberts, P.A., Xu, S., Fatokun, C. and Close, T.J. (2018) Identification of QTL controlling domestication-related traits in cowpea (*Vigna unguiculata* L. Walp). Scientific Reports, 8, 6261.

Lonardi, S., Muñoz-Amatriaín, M., Liang, Q., Shu, S., Wanamaker, S.I., Lo, S., Tanskanen, J., Schulman, A.H., Zhu, T., Luo, M.C., Alhakami, H., Ounit, R., Hasan, A.M., Verdier, J., Roberts, P.A., Santos, J.R.P., Ndeve, A., Doležel, J., Vrána, J., Hokin, S.A., Farmer, A.D., Cannon, S.B. and Close, T.J. (2019) The genome of cowpea (Vigna unguiculata [L.] Walp.). biorxiv, 518969.

Lush, W.M. and Evans, L.T. (1981) The domestication and improvement of cowpeas (*Vigna unguiculatea* (L.) Walp.). Euphytica, 30, 579–587.

Mitsuda, N. and Ohme-Takagi, M. (2008) NAC transcription factors NST1 and NST3 regulate pod shattering in a partially redundant manner by promoting secondary wall formation after the establishment of tissue identity. Plant Journal, 56, 768–778.

Murgia, M.L., Attene, G., Rodriguez, M., Bitocchi, E., Bellucci, E., Fois, D., Nanni, L., Gioia, T., Albani, D.M., Papa, R. and Rau, D. (2017) A Comprehensive Phenotypic Investigation of the “Pod-Shattering Syndrome” in Common Bean. Frontiers in Plant Science, 8, 251.

Parker, T.A., Miery Teran, J.C.B., Palkovic, A., Jernstedt, J. and Gepts, P. (2019) Genetic control of pod dehiscence in domesticated common bean: Associations with range expansion and local aridity conditions. biorxiv, 517516.

Philbrook, B. and Oplinger, E.S. (1989) Soybean field losses as influenced by harvest delays. Agronomy Journal, 81, 251–258.

Pormar, F., Merino, F. and Barceló, R. (2002) O-4-linked coniferyl and sinapyl alehydes in lignifying cell walls are the main targets of the Wiesner (phloroglucinol-HCl) reaction. Protoplasma, 220, 17–28.

Rau, D., Murgia, M.L., Rodriguez, M., Bitocchi, E., Bellucci, E., Fois, D., Albani, D., Nanni, L., Gioia, T., Santo, D., Marcolungo, L., Delledonne, M., Attene, G. and Papa, R. (2019) Genomic dissection of pod shattering in common bean: mutations at non-orthologous loci at the basis of convergent phenotypic evolution under domestication of leguminous species. Plant Journal, 97, 693–714.

Sakai, H., Naito, K., Ogiso-Tanaka, E., Takahashi, Y., Iseki, K., Muto, C., Satou, K., Teruya, K., Shiroma, A., Shimoji, M., Hirano, T., Itoh, T., Kaga, A. and Tomooka, N. (2015) The power of single molecule real-time sequencing technology in the de novo assembly of a eukaryotic genome. Scientific Reports, 5, 16780.

Sakai, H., Naito, K., Takahashi, Y., Sato, T., Yamamoto, T., Muto, I., Itoh, T. and Tomooka, N. (2016) The *Vigna* Genome Server, ‘*Vig*GS’: A Genomic Knowledge Base of the Genus *Vigna* Based on High-Quality, Annotated Genome Sequence of the Azuki Bean, *Vigna angularis* (Willd.) Ohwi & Ohashi. Plant and Cell Physiology, 57, e2.

Smit, A.F.A., Hubley, R. and Green, P. (2013-2015) RepeatMasker Open-4.0. (http://www.repeatmasker.org).

Smit, A.F.A. and Hubley, R. (2008-2015) RepeatModeler Open-1.0. (http://www.repeatmasker.org).

Spence, J., Vercher, Y., Gates, P. and Harris, N. (1996) ‘Pod shatter’ in *Arabidopsis thaliana*, Brassica napus and B. juncea. Journal of Microscopy, 181, 195–203.

Stanke, M., Diekhans, M., Baetsch, R. and Haussler, D. (2008) Using native and syntenically mapped cDNA alignments to improve de novo gene finding. Bioinformatics, 24, 637–664.

Suanum, W., Somta, P., Kongjaimun, A., Yimram, T., Kaga, A., Tomooka, N., Takahashi, Y. and Srinives, P. (2016) Co-localization of QTLs for pod Fiber content and pod shattering in F 2 and backcross populations between yardlong bean and wild cowpea. Molecular Breeding, 36, 80.

Untergasser, A., Cutcultache, I., Koressaar, T., Faircloth, B.C., Remm, M. and Rozen, S.G. (2012) Primer3—new capabilities and interfaces. Bioinformatics, 40, e115.

Waterhouse, R.M., Seppey, M., Simão, F.A., Manni, M., Ioannidis, P., Klioutchnikov, G., Kriventseva, E.V. and Zdobnov, E.M. (2017) BUSCO Applications from Quality Assessments to Gene Prediction and Phylogenomics. Molecular Biology and Evolution, 35, 543–548.

Yang, C., Xu, Z., Song, J., Conner, K., Vizcay Barrena, G. and Wilson, Z.A. (2007) Arabidopsis MYB26/MALE STERILE35 regulates secondary thickening in the endothecium and is essential for anther dehiscence. Plant Cell, 19, 534–548.

Yang, C., Song, J., Ferguson, A.C., Klisch, D., Simpson, K., Mo, R., Taylor, B., Mitsuda, N. and Wilson, Z.A. (2017) Transcription Factor MYB26 Is Key to Spatial Specificity in Anther Secondary Thickening Formation. Plant Physiology, 175, 333–350.

Zhao, Q. and Dixon, R.A. (2011) Transcriptional networks for lignin biosynthesis: more complex than we thought? Trends in Plant Science, 16, 227–233.

