## SupplementalFigures for "Genetic factor for twisting legume pods identified by fine-mapping of shattering-related traits in azuki bean and yard-long bean"

### Slide 1
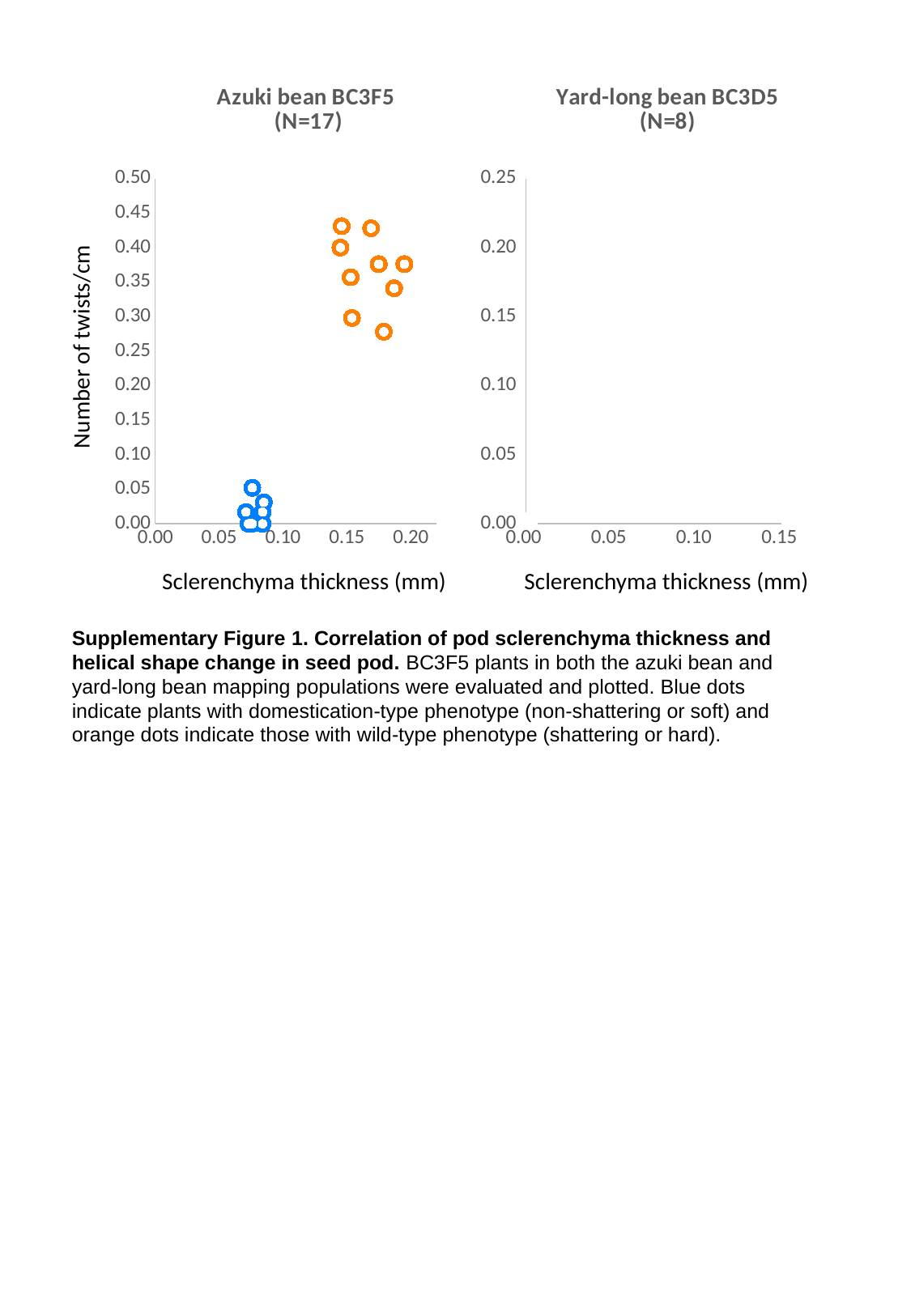

#### Chart: Azuki bean BC3F5
(N=17)
| Category |
|---|
#### Chart: Yard-long bean BC3D5 (N=8)
| Category | | | |
|---|---|---|---|Number of twists/cm
Sclerenchyma thickness (mm)
Sclerenchyma thickness (mm)
Supplementary Figure 1. Correlation of pod sclerenchyma thickness and helical shape change in seed pod. BC3F5 plants in both the azuki bean and yard-long bean mapping populations were evaluated and plotted. Blue dots indicate plants with domestication-type phenotype (non-shattering or soft) and orange dots indicate those with wild-type phenotype (shattering or hard).

### Slide 2
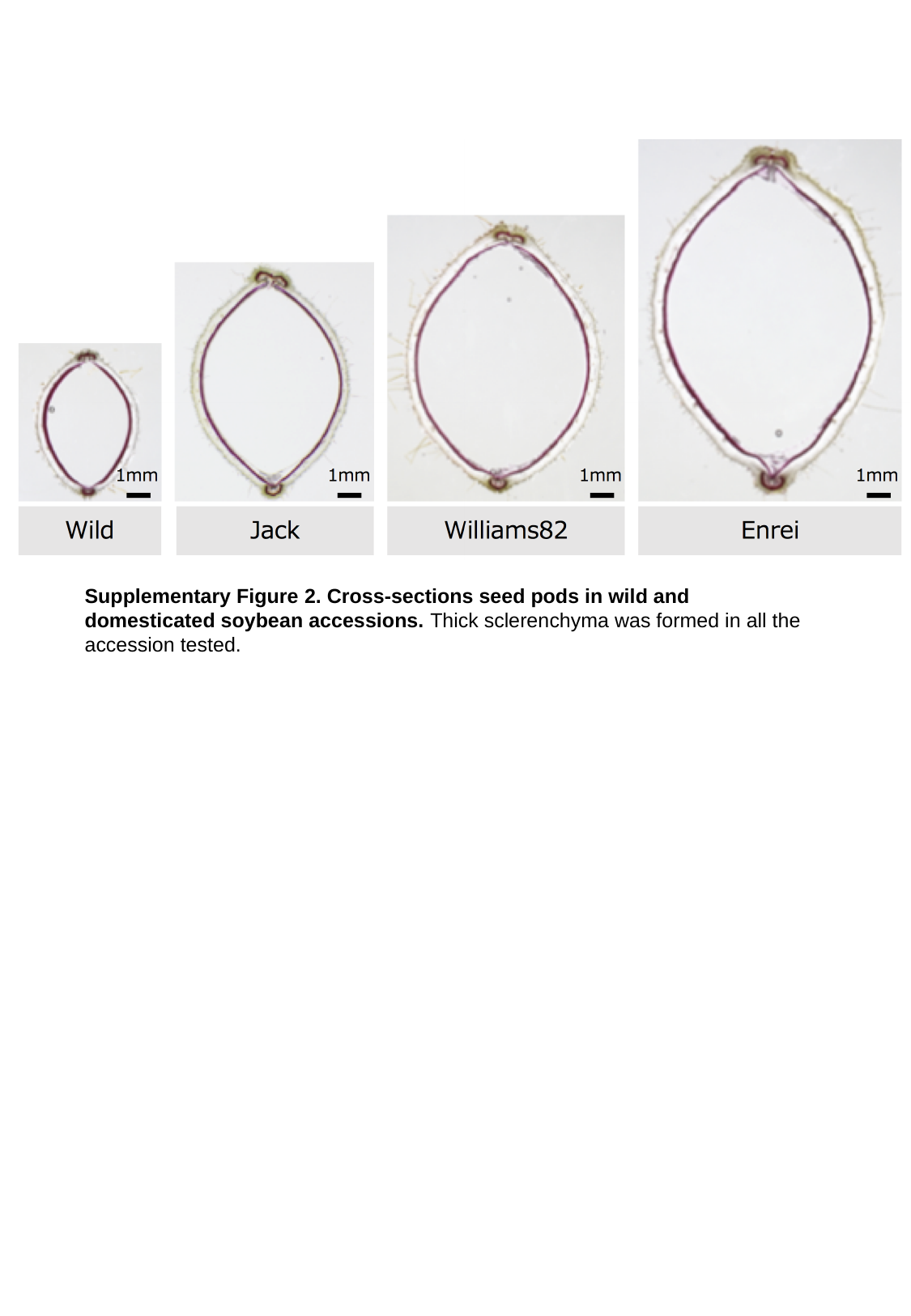

Supplementary Figure 2. Cross-sections seed pods in wild and domesticated soybean accessions. Thick sclerenchyma was formed in all the accession tested.

### Slide 3
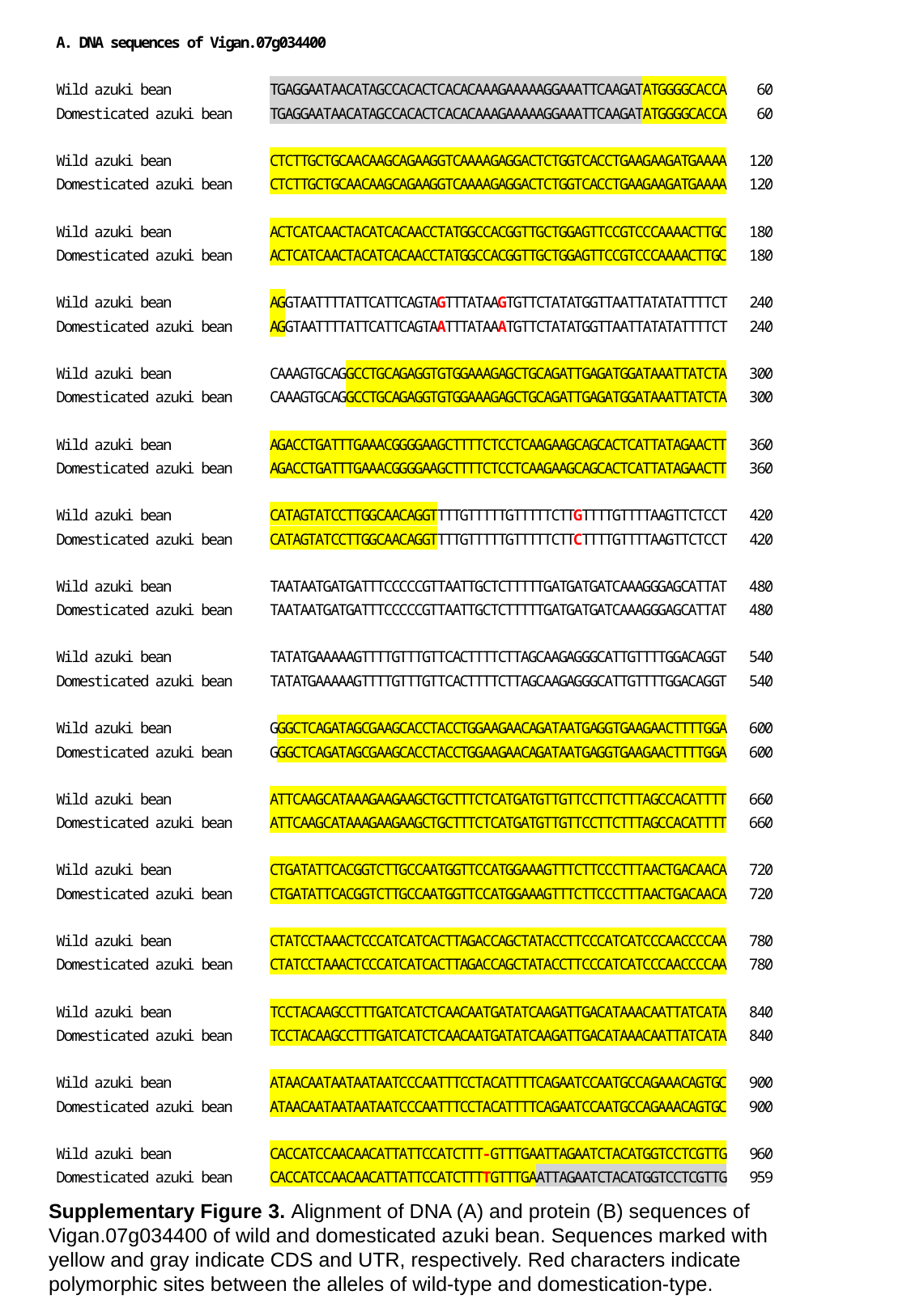

Supplementary Figure 3. Alignment of DNA (A) and protein (B) sequences of Vigan.07g034400 of wild and domesticated azuki bean. Sequences marked with yellow and gray indicate CDS and UTR, respectively. Red characters indicate polymorphic sites between the alleles of wild-type and domestication-type.

### Slide 4
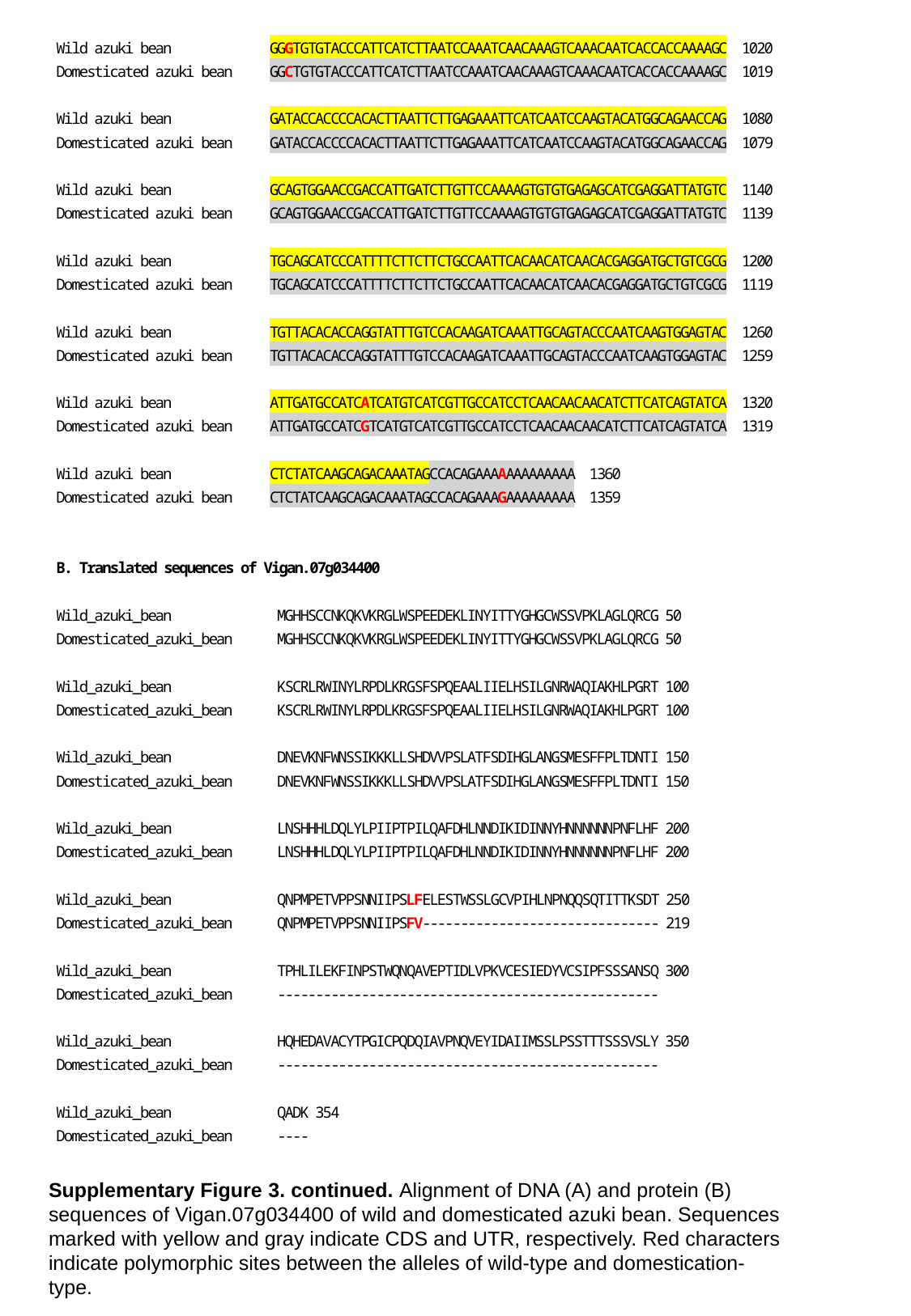

Supplementary Figure 3. continued. Alignment of DNA (A) and protein (B) sequences of Vigan.07g034400 of wild and domesticated azuki bean. Sequences marked with yellow and gray indicate CDS and UTR, respectively. Red characters indicate polymorphic sites between the alleles of wild-type and domestication-type.

### Slide 5
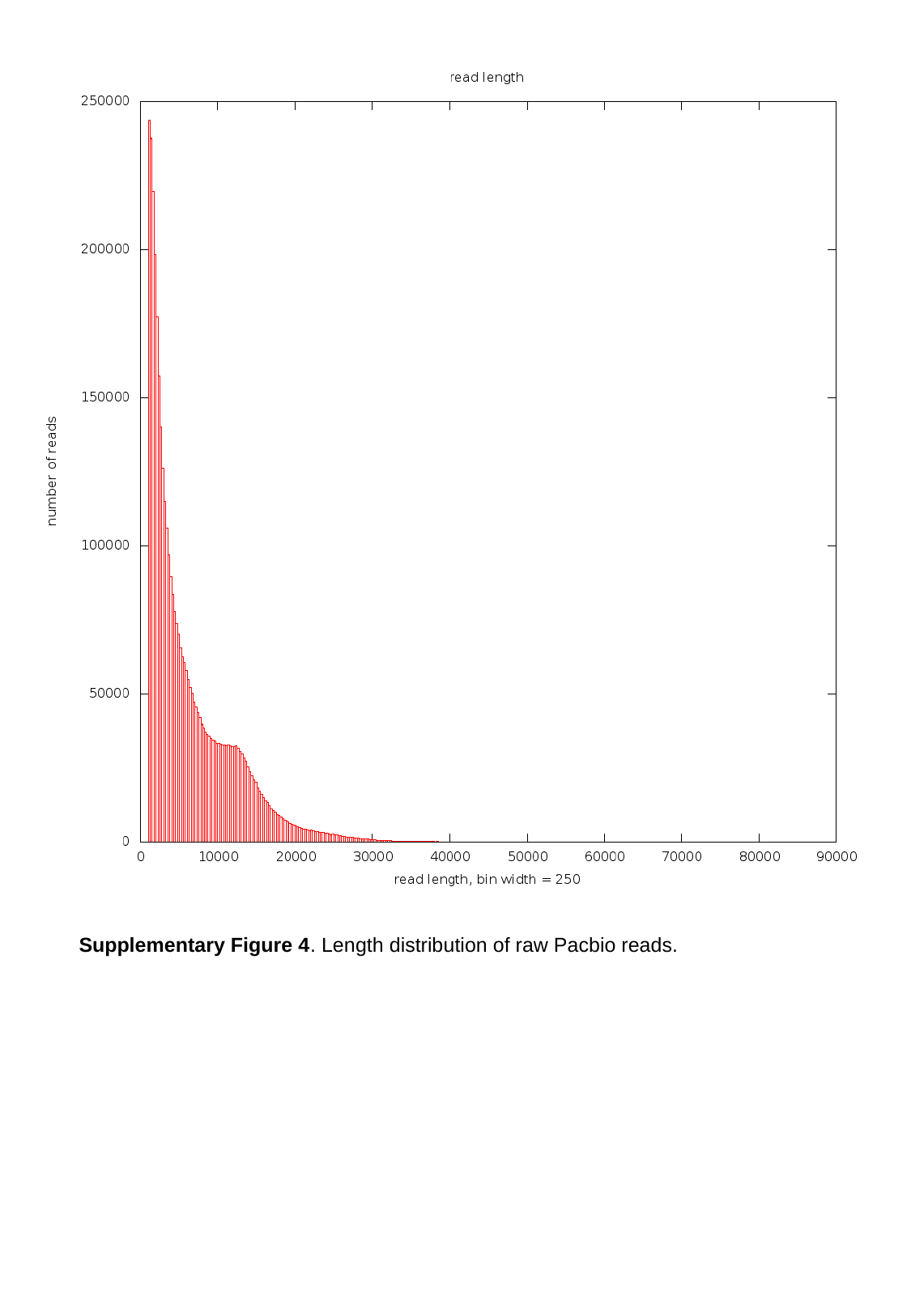

Supplementary Figure 4. Length distribution of raw Pacbio reads.

### Slide 6
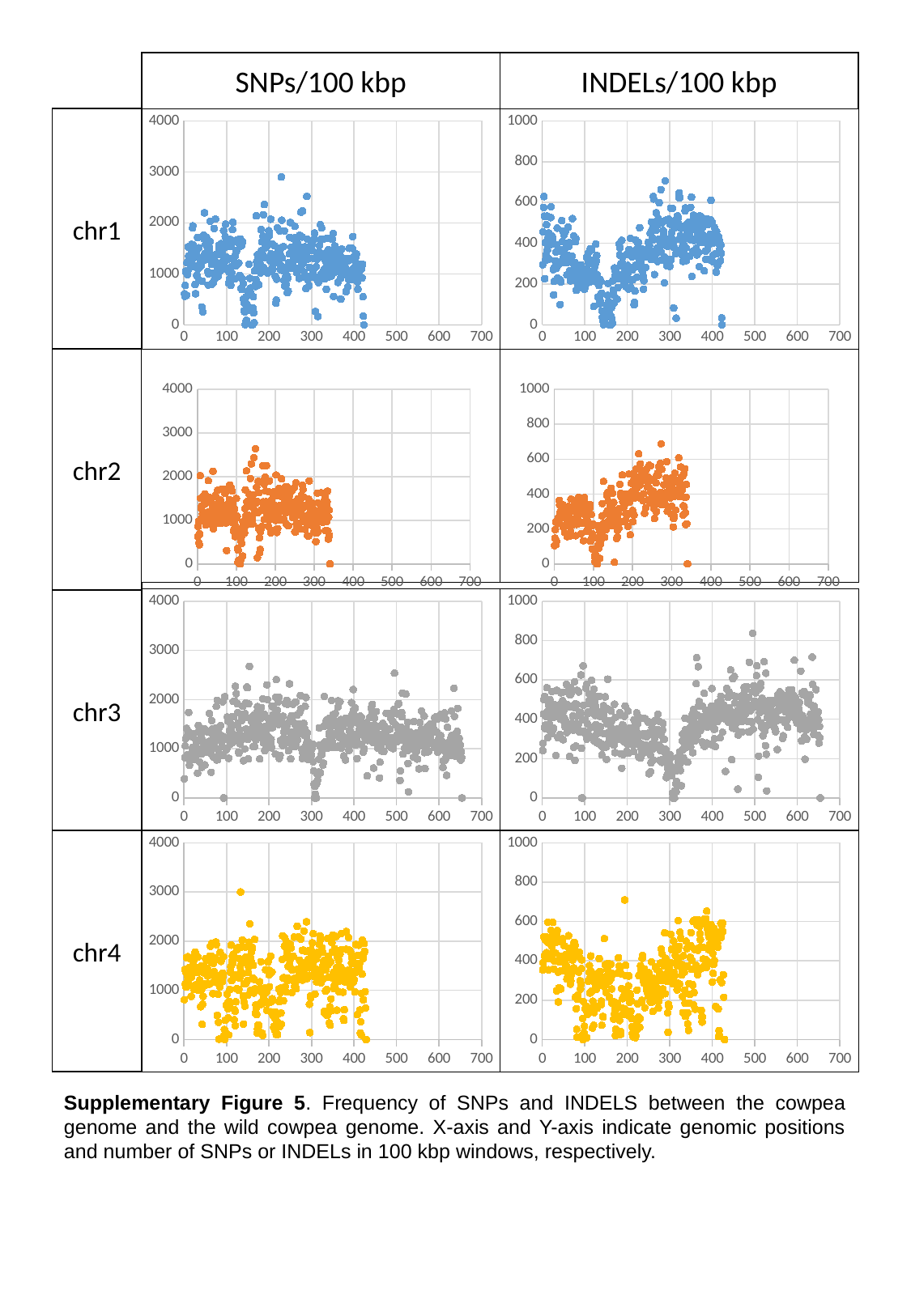

SNPs/100 kbp
INDELs/100 kbp
chr1
#### Chart
| Category |
|---|
#### Chart
| Category | |
|---|---|chr2
#### Chart
| Category |
|---|
#### Chart
| Category |
|---|
#### Chart
| Category |
|---|
#### Chart
| Category | |
|---|---|chr3
chr4
#### Chart
| Category |
|---|
#### Chart
| Category | |
|---|---|Supplementary Figure 5. Frequency of SNPs and INDELS between the cowpea genome and the wild cowpea genome. X-axis and Y-axis indicate genomic positions and number of SNPs or INDELs in 100 kbp windows, respectively.

### Slide 7
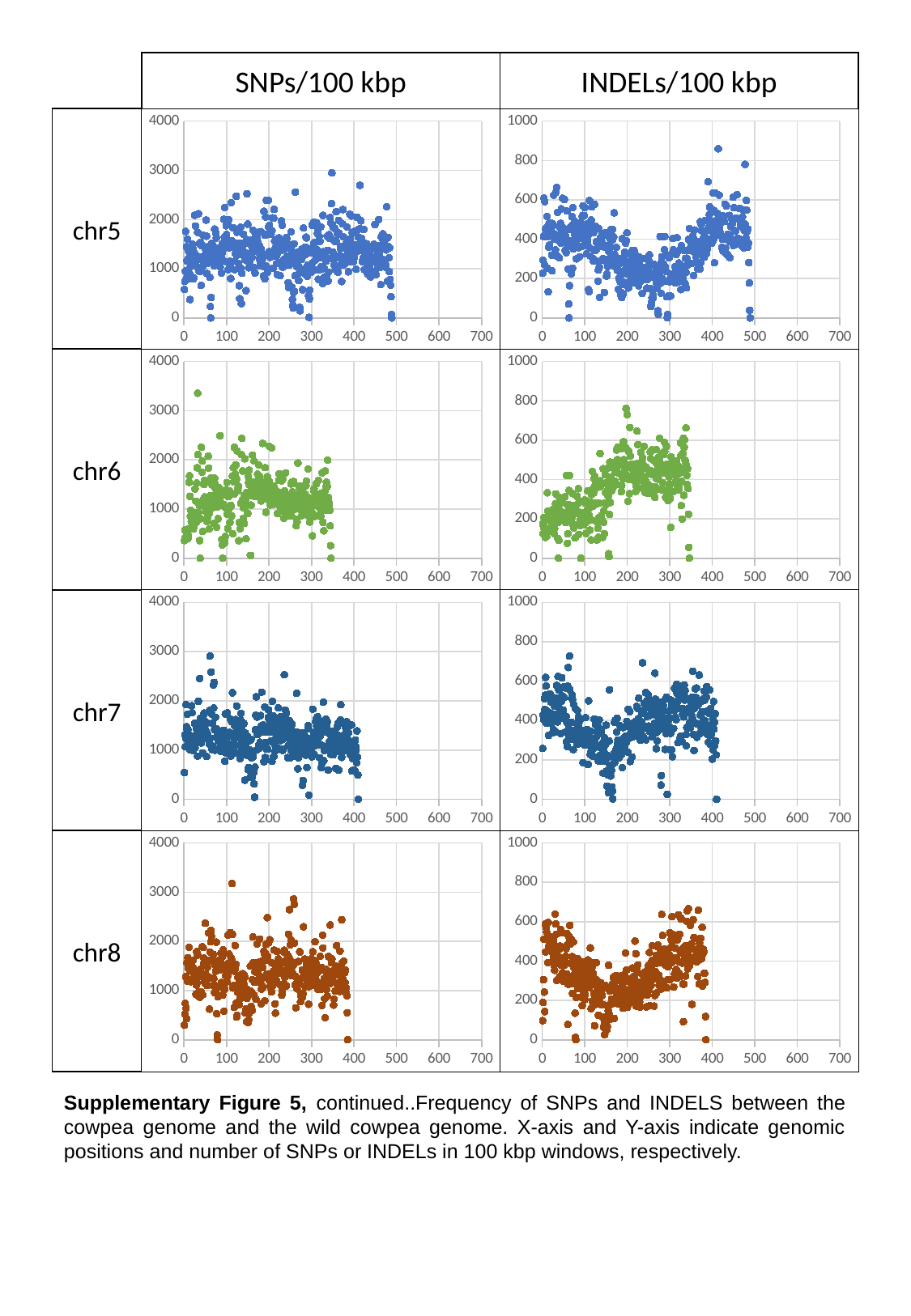

SNPs/100 kbp
INDELs/100 kbp
chr5
#### Chart
| Category |
|---|
#### Chart
| Category | |
|---|---|chr6
#### Chart
| Category |
|---|
#### Chart
| Category | |
|---|---|chr7
#### Chart
| Category |
|---|
#### Chart
| Category | |
|---|---|chr8
#### Chart
| Category |
|---|
#### Chart
| Category | |
|---|---|Supplementary Figure 5, continued..Frequency of SNPs and INDELS between the cowpea genome and the wild cowpea genome. X-axis and Y-axis indicate genomic positions and number of SNPs or INDELs in 100 kbp windows, respectively.

### Slide 8
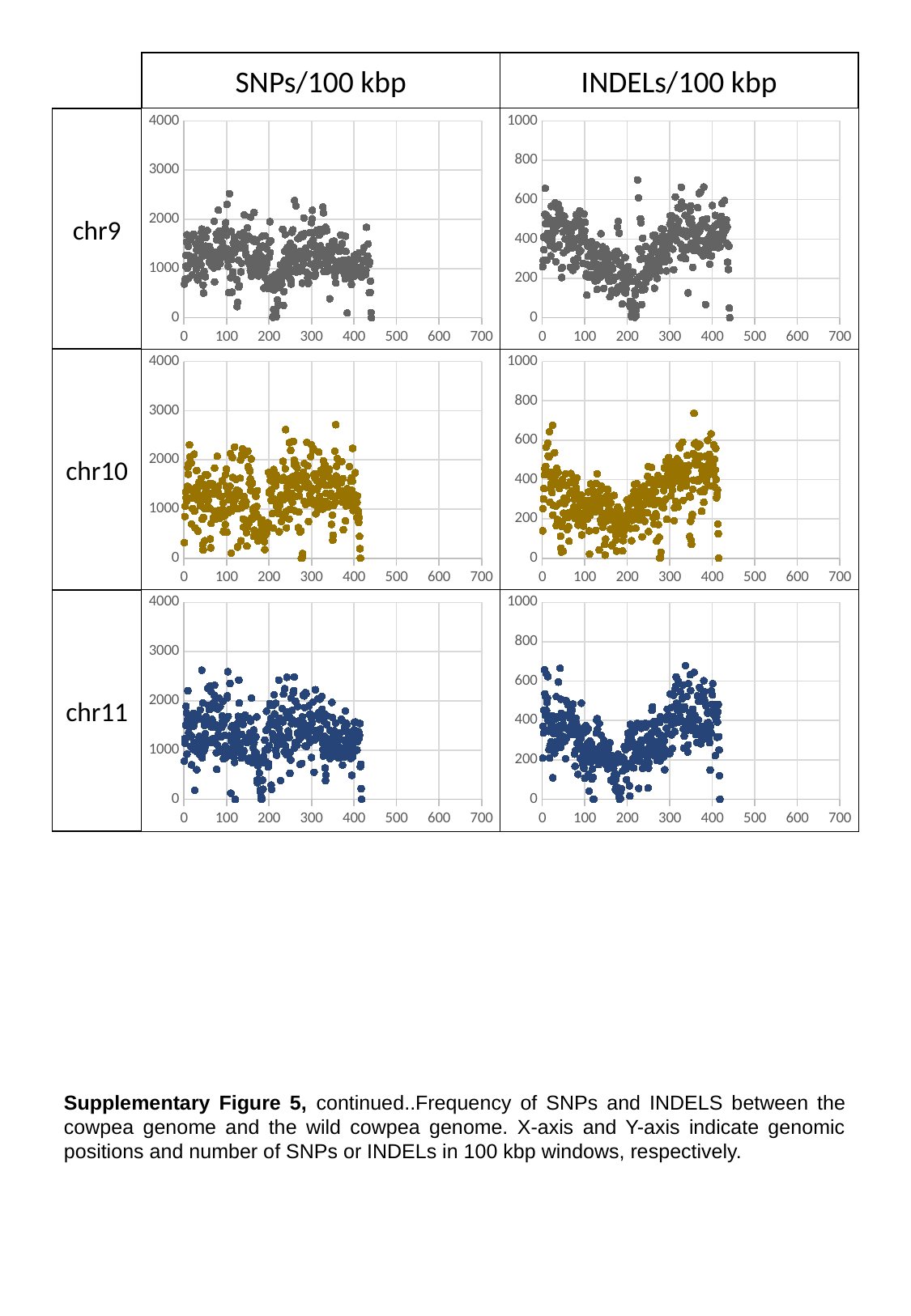

SNPs/100 kbp
INDELs/100 kbp
chr9
#### Chart
| Category |
|---|
#### Chart
| Category | |
|---|---|chr10
#### Chart
| Category |
|---|
#### Chart
| Category | |
|---|---|chr11
#### Chart
| Category |
|---|
#### Chart
| Category | |
|---|---|Supplementary Figure 5, continued..Frequency of SNPs and INDELS between the cowpea genome and the wild cowpea genome. X-axis and Y-axis indicate genomic positions and number of SNPs or INDELs in 100 kbp windows, respectively.

### Slide 9
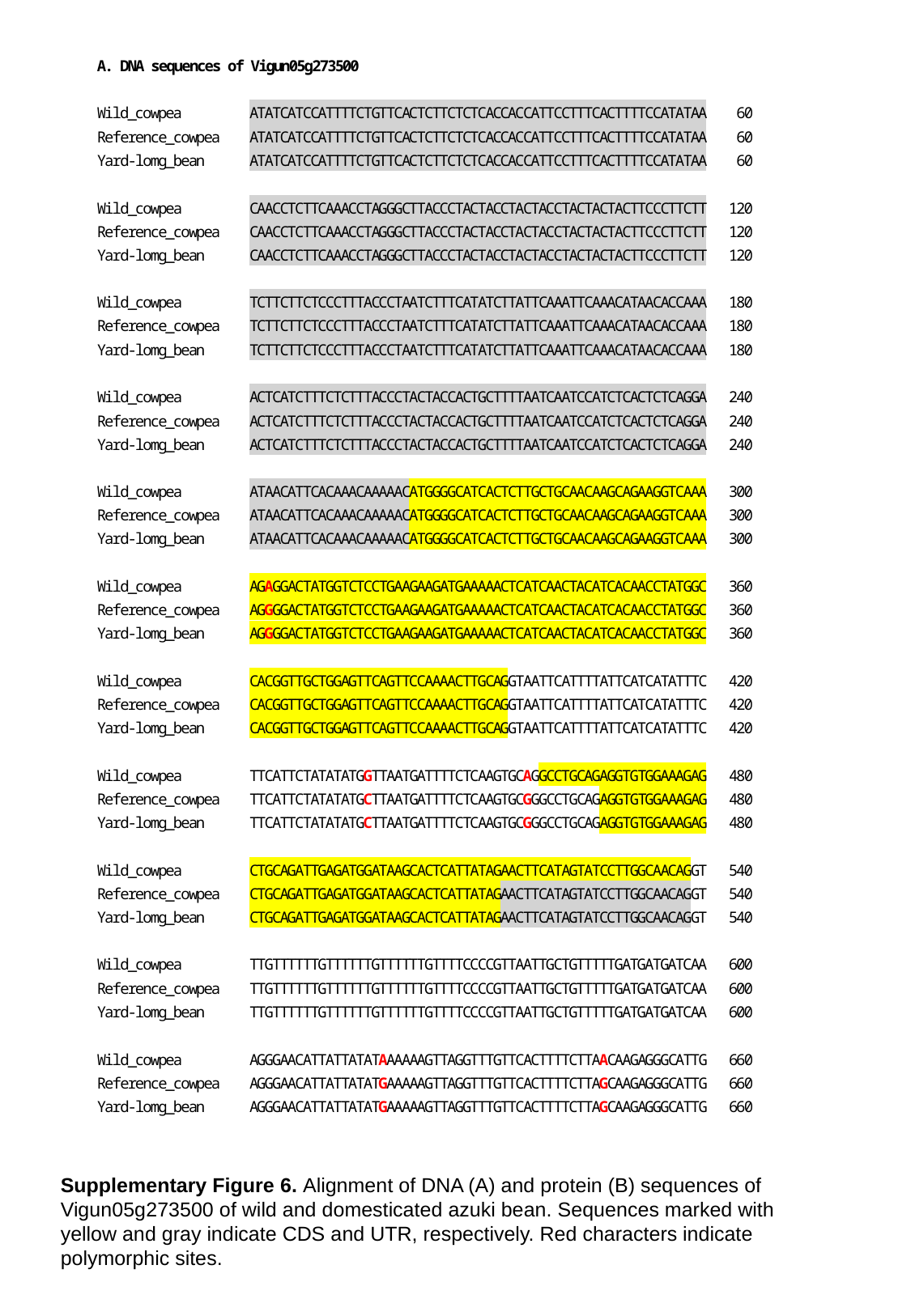

Supplementary Figure 6. Alignment of DNA (A) and protein (B) sequences of Vigun05g273500 of wild and domesticated azuki bean. Sequences marked with yellow and gray indicate CDS and UTR, respectively. Red characters indicate polymorphic sites.

### Slide 10
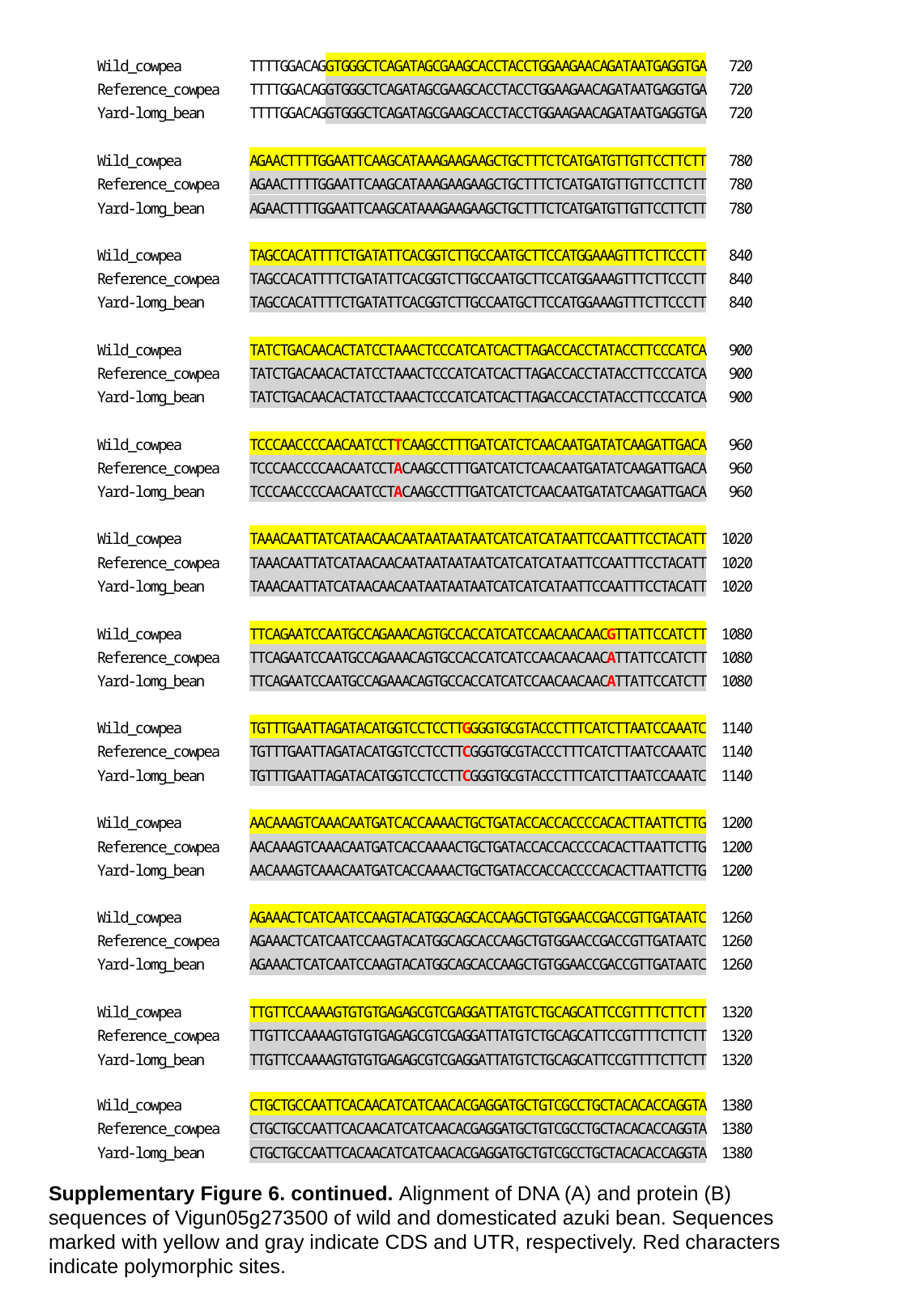

Supplementary Figure 6. continued. Alignment of DNA (A) and protein (B) sequences of Vigun05g273500 of wild and domesticated azuki bean. Sequences marked with yellow and gray indicate CDS and UTR, respectively. Red characters indicate polymorphic sites.

### Slide 11
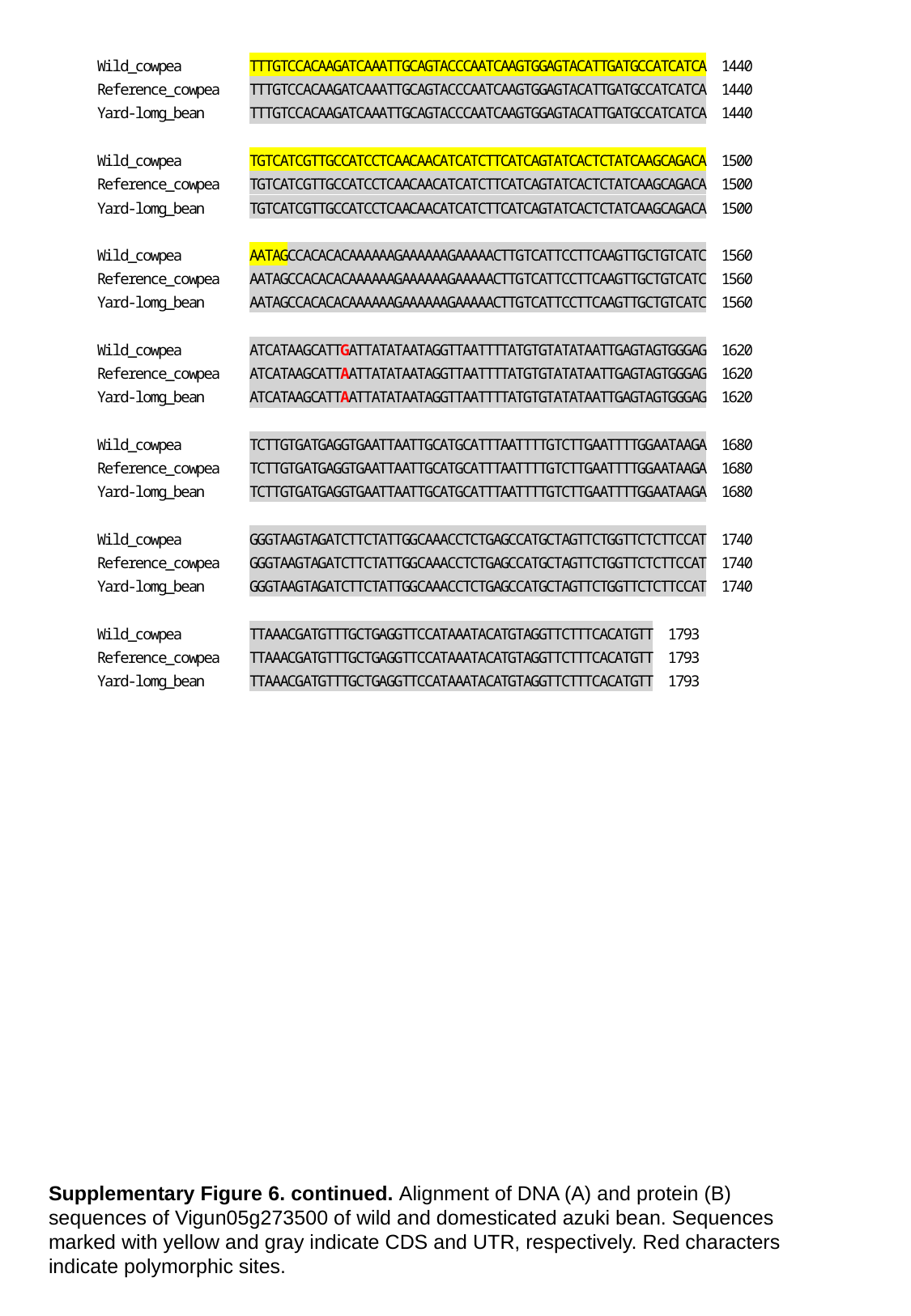

Supplementary Figure 6. continued. Alignment of DNA (A) and protein (B) sequences of Vigun05g273500 of wild and domesticated azuki bean. Sequences marked with yellow and gray indicate CDS and UTR, respectively. Red characters indicate polymorphic sites.

### Slide 12
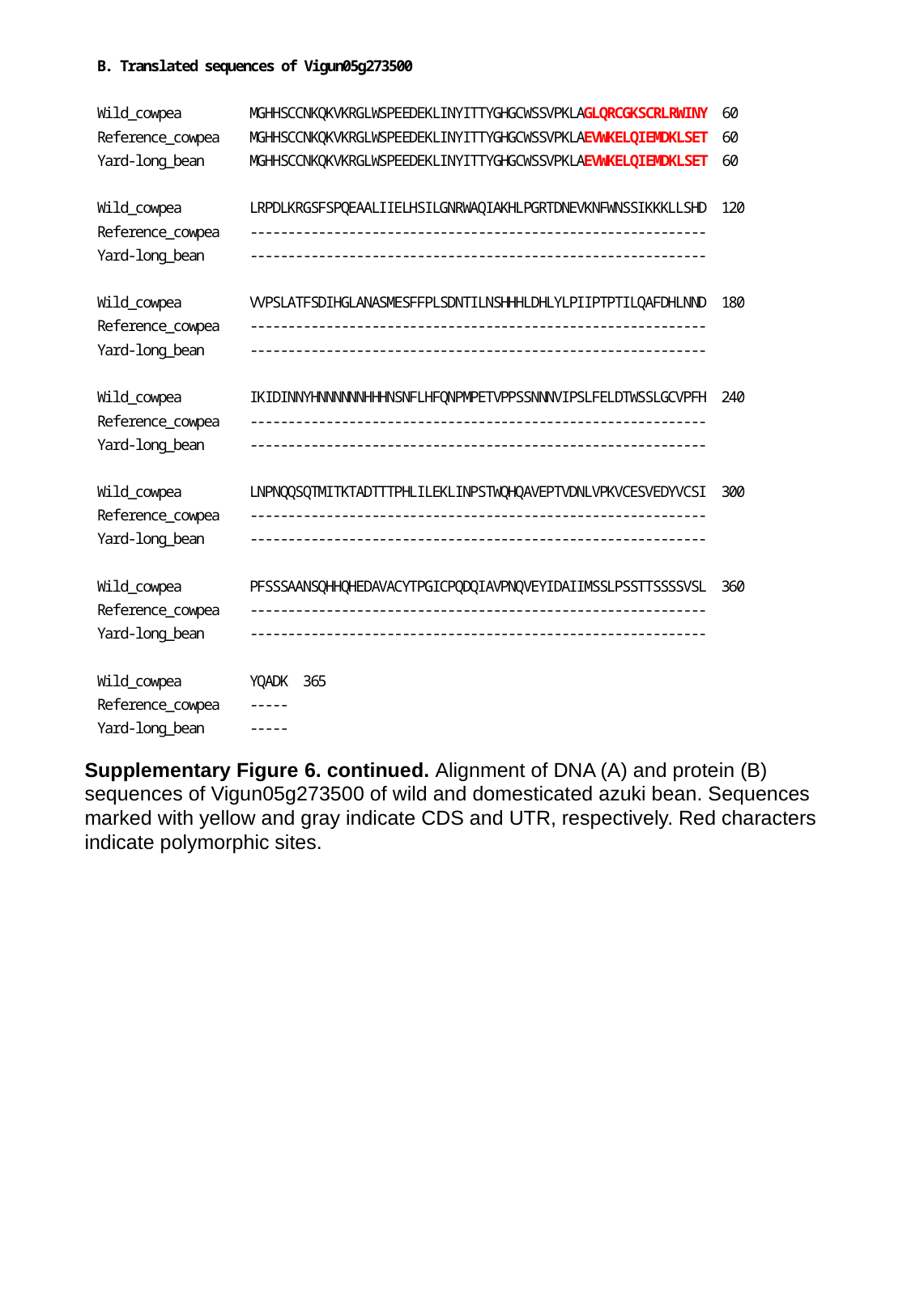

Supplementary Figure 6. continued. Alignment of DNA (A) and protein (B) sequences of Vigun05g273500 of wild and domesticated azuki bean. Sequences marked with yellow and gray indicate CDS and UTR, respectively. Red characters indicate polymorphic sites.

### Slide 13
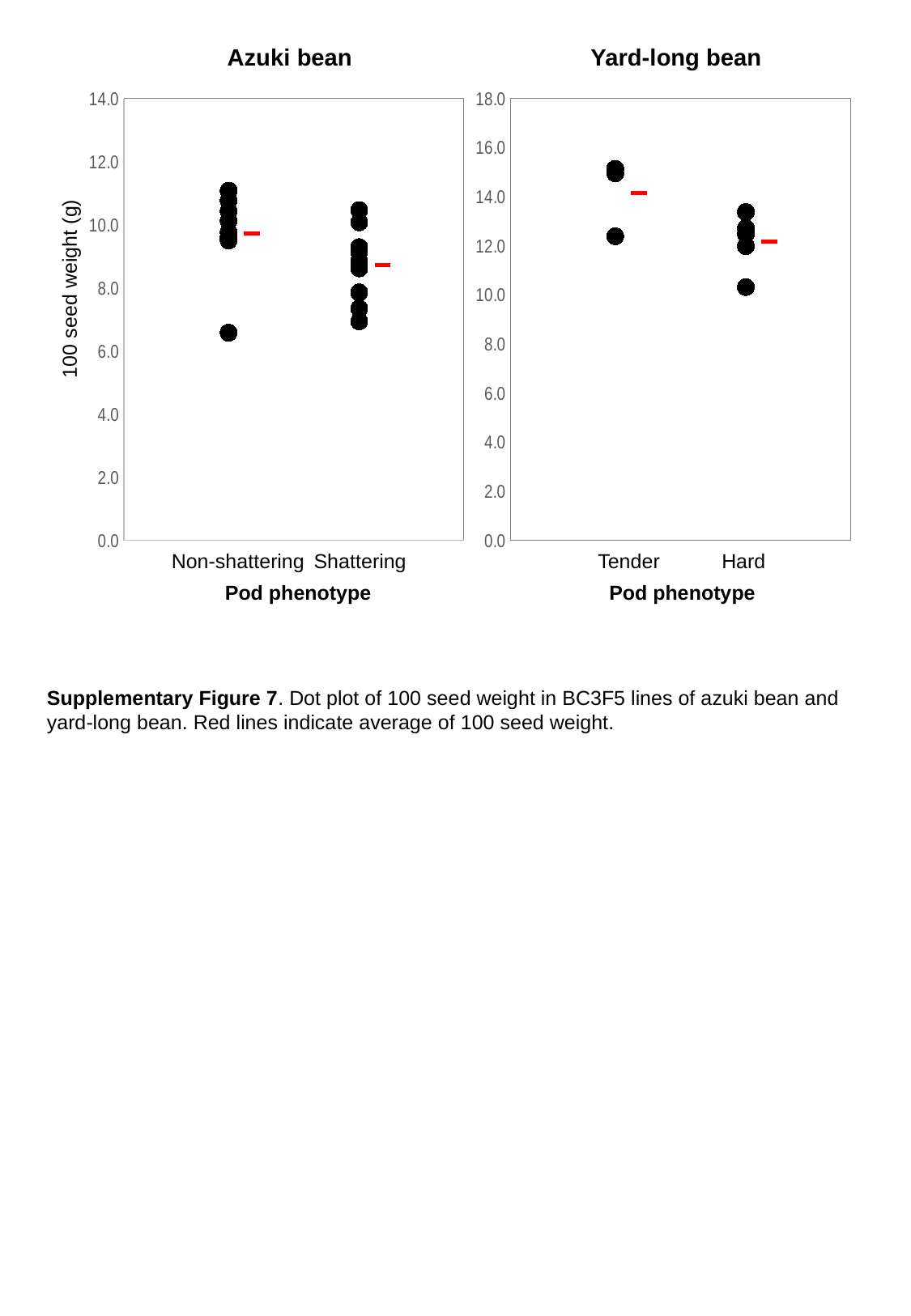

Azuki bean
Yard-long bean
#### Chart
| Category |
|---|
#### Chart
| Category | | |
|---|---|---|100 seed weight (g)
Non-shattering
Shattering
Tender
Hard
Pod phenotype
Pod phenotype
Supplementary Figure 7. Dot plot of 100 seed weight in BC3F5 lines of azuki bean and yard-long bean. Red lines indicate average of 100 seed weight.

### Slide 14
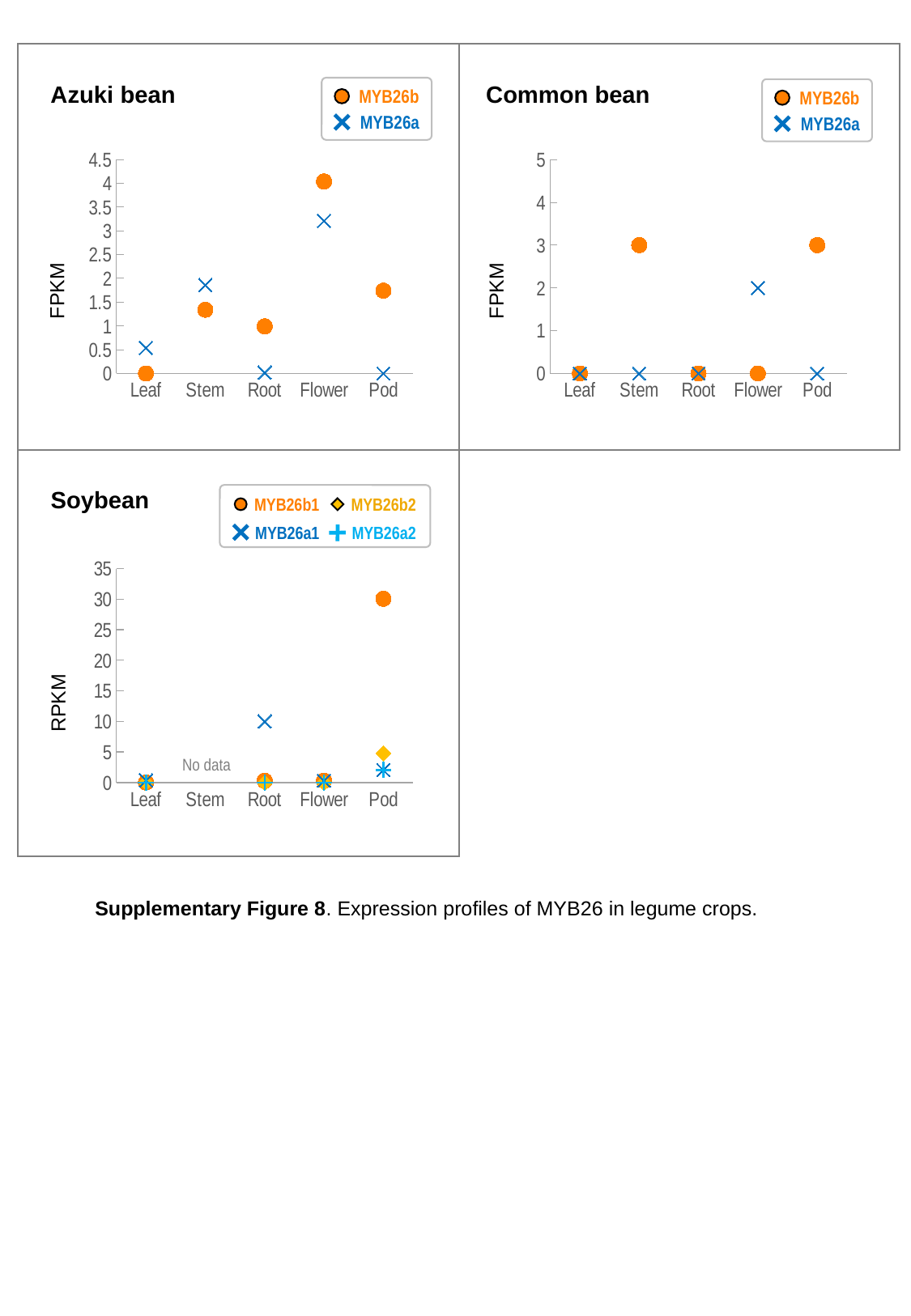

MYB26b
MYB26a
MYB26b
MYB26a
Azuki bean
Common bean
#### Chart
| Category | VaMYBa | VaMYBb |
|---|---|---|
| Leaf | 0.0 | 0.54 |
| Stem | 1.34 | 1.86 |
| Root | 0.99 | 0.02 |
| Flower | 4.04 | 3.21 |
| Pod | 1.74 | 0.0 |
#### Chart
| Category | PvMYBa | PvMYBb |
|---|---|---|
| Leaf | 0.0 | 0.0 |
| Stem | 3.0 | 0.0 |
| Root | 0.0 | 0.0 |
| Flower | 0.0 | 2.0 |
| Pod | 3.0 | 0.0 |FPKM
FPKM
Soybean
MYB26b1
MYB26b2
MYB26a1
MYB26a2
#### Chart
| Category | GmMYBa1 | GmMYBa2 | GmMYBb1 | GmMYBb2 |
|---|---|---|---|---|
| Leaf | 0.0 | 0.0 | 0.35 | 0.0 |
| Stem | None | None | None | None |
| Root | 0.26 | 0.0 | 10.0 | 0.0 |
| Flower | 0.29 | 0.0 | 0.29 | 0.0 |
| Pod | 30.06 | 4.78 | 2.04 | 2.04 |RPKM
No data
Supplementary Figure 8. Expression profiles of MYB26 in legume crops.
